# Multidimensional profiling of drug-treated cells by Imaging Mass Cytometry

**DOI:** 10.1101/549592

**Authors:** Alexandre Bouzekri, Amanda Esch, Olga Ornatsky

## Abstract

In pharmaceutical research, high-content screening is an integral part of lead candidate development. Drug response *in vitro* by examining over 40 parameters, including biomarkers, signaling molecules, cell morphological changes, proliferation indices and toxicity in a single sample, could significantly enhance discovery of new therapeutics. As a proof of concept, we present here a workflow for multidimensional Imaging Mass Cytometry™ (IMC™) and data processing with open source computational tools. CellProfiler was used to identify single cells through establishing cellular boundaries, followed by histoCAT™ (histology topography cytometry analysis toolbox) for extracting single-cell quantitative information visualized as t-SNE plots and heatmaps. Human breast cancer-derived cell lines SKBR3, HCC1143 and MCF-7 were screened for expression of cellular markers to generate digital images with a resolution comparable to conventional fluorescence microscopy. Predicted pharmacodynamic effects were measured in MCF-7 cells dosed with three target-specific compounds: growth stimulatory EGF, microtubule depolymerization agent nocodazole and genotoxic chemotherapeutic drug etoposide. We show strong pairwise correlation between nuclear markers pHistone3^S28^, Ki-67 and p4E-BP1^T37/T46^ in classified mitotic cells and anti-correlation with cell surface markers. Our study demonstrates that IMC data expand the number of measured parameters in single cells and brings higher-dimension analysis to the field of cell-based screening in early lead compound discovery.

## Introduction

Discovery of new treatments in oncology research relies extensively on the use of human-derived cell culture models [1]. High-content cell-based screens are widely applied in pharmaceutical drug development to prioritize lead molecules for animal testing [2]. These assays rely on the use of primary and cancer cell lines and mostly monitor cytotoxicity and proliferation. With the advent of sophisticated genomics and proteomics technologies for soluble proteins and mass cytometry for single cells, it is becoming possible to increase the multidimensionality of *in vitro* screens. Databases of genotypic and phenotypic profiles [3, 4, 5] across cancer drug panels have provided a comparative analysis between *in vitro* studies and clinical therapeutic responses *in vivo* [6]. One of these datasets developed and shared by the National Cancer Institute 60 (NCI-60) in the late 1980s was the first *in vitro* discovery screening tool for pharmacologic compounds inducing growth inhibition across 60 cancer cell lines [7]. Over the years several concerns were raised regarding the use of *in vitro* established human tumor-derived cell lines for drug testing. Concerns included genetic instability and dedifferentiation. However, the study of genetic mutations arising in immortalized cells remained a field of interest in drug discovery, providing knowledge on the dysregulation of cellular signaling pathways and the effects small-molecule inhibitors have on human tumor cell lines [8]. High-content imaging of cancer cell lines in response to drug treatment is a standard assay applied in preclinical studies for identification of different mechanisms of drug action [9, 10, 11, 12]. The main criteria in every morphophenotypic screen for early drug discovery is the selection of biomarkers and detection modalities (antibodies, chemical and enzymatic probes, reporter detection tags). In the field of fluorescent cellular imaging microscopy, biomarker analysis in single cells is limited to 4–6 due to the overlapping spectra of fluorescent dyes [13]. Repetitive rounds of staining using the same biological sample are required to achieve multiplexing [14]. New imaging technologies are needed to significantly increase multiplicity in a single sample preparation experiment and perform replicate analyses [15]. High-content imaging assays applied to screening drug perturbations in heterogeneous cancer cell models could uncover additional modes of drug action and biomarkers for clinical trials [16].

Imaging Mass Cytometry is an emerging and transformative technique in the field of digital histopathology applied to complex tissue sections [17, 18, 19]. The Hyperion™ Imaging System (Fluidigm^®^) can measure up to 40 parameters simultaneously in formalin-fixed, paraffin-embedded (FFPE) and frozen human tissue sections with subcellular resolution [20]. Sample preparation is very similar to standard immunohistochemical protocols [17], where the tissue section on a slide is first deparaffinized and treated with an antigen retrieval buffer for antigen epitope exposure in the case of FFPE, then stained once with a mixture of isotopically pure metal-labeled antibodies specific to structural and cell type biomarkers. After drying and inserting the slide into the ablation chamber, regions of interest (ROIs) are chosen and recorded by a camera integrated with the Hyperion Tissue Imager. These ROIs are then ablated in 1 μm steps as the slide moves under the laser. Each laser pulse vaporizes a 1 μm^2^ area of the tissue and generates a plume of particles. The resulting particles are carried by a stream of helium/argon mixture into a CyTOF^®^ instrument, an inductively coupled plasma time-of-flight mass cytometer, described elsewhere [18]. Data are stored as raw binary files of ion counts for each mass channel (parameter) per pixel. These files can be converted into a stack of single-parameter matrices (grayscale images), which are used to create merged multiparametric pseudocolor images and perform quantitative pixel- or segmentation-based analysis. Bodenmiller et al. are developing multivariate computational tools to visualize and analyze multiplexed images of human tissue sections generated by IMC [21]. However, there is no single software package or analysis workflow that could currently be applied to answer specific biological questions.

In this proof-of-principle study, we set out to develop a comprehensive workflow for IMC data analysis based on recent advances in imaging algorithmic methods to visualize and measure multiple biomarkers in model cell lines cultured in chamber slides **(Fig. 1)**. Several human breast cancer-derived cell lines, SKBR3, HCC1143 and MCF-7, were screened for expression of surface and intracellular markers. Predicted pharmacodynamic effects were studied in MCF-7 cells dosed with three target-specific compounds: growth stimulatory epidermal growth factor (EGF), microtubule depolymerization agent nocodazole and genotoxic chemotherapeutic drug etoposide [22, 23]. Our method analysis workflow demonstrates the high multiplexing capability of IMC for *in vitro* research and offers quantitation of multiple biomarkers at subcellular resolution. Future improvements of the technology toward a higher acquisition speed of multiple samples will expand IMC applications as an imaging platform for *in vitro* cell-based drug screening.

**Figure 1.**
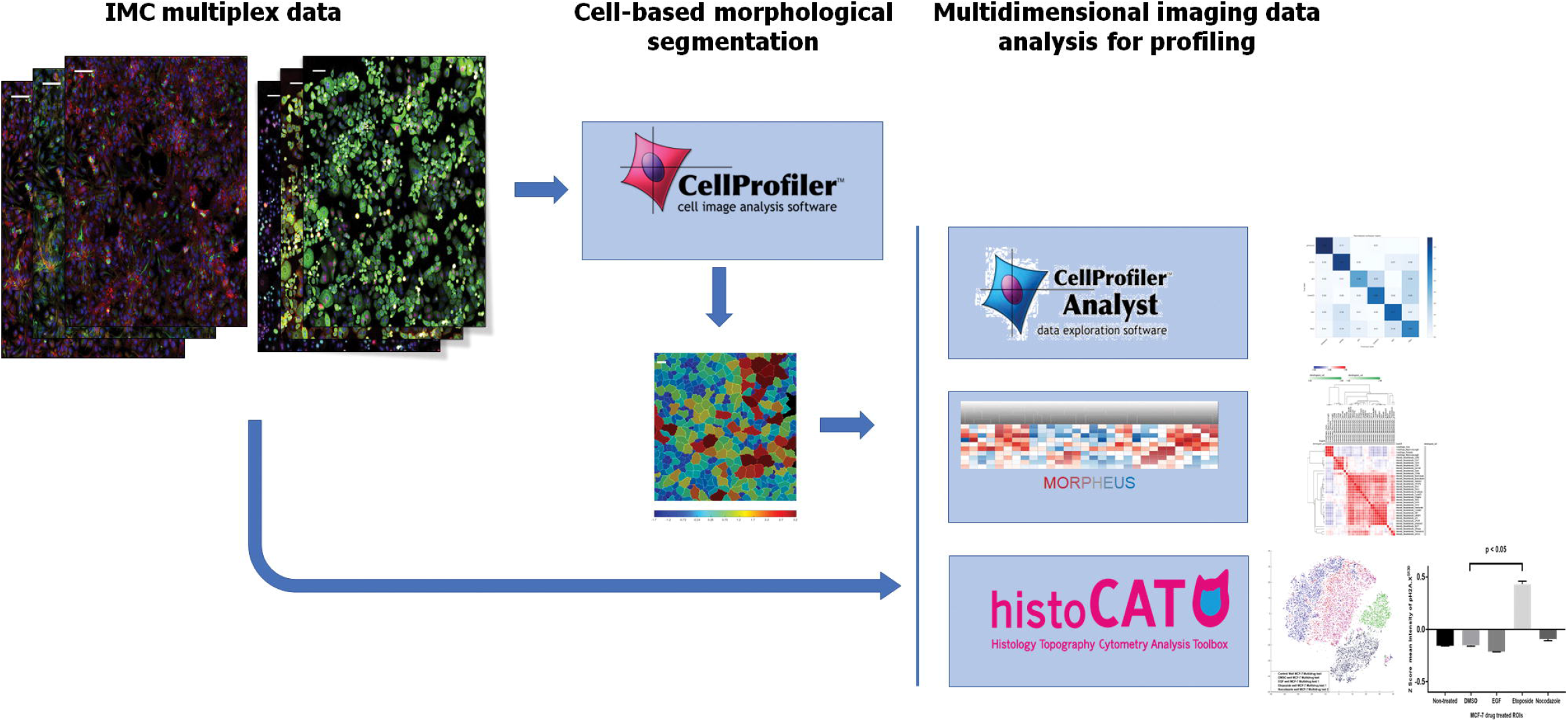
Image-based IMC data analysis workflow with open source software.

## Material and Methods

### Cell culture

SKBR3 (HTB-30™), HCC1143 (CRL-2321™) and MCF-7 (HTB-22™) were purchased, tested for mycoplasma contamination and authenticated with short tandem repeat DNA profiling by American Tissue Culture Collection (ATCC^®^). Cells were cultured within 15 passages in their corresponding growth media supplemented with 10% bovine serum (HyClone™ Cosmic Calf™ Serum, Cat. No. SH30087.04) and penicillin/streptomycin (Thermo Fisher Scientific Cat. No. 15140-122): McCoy’s 5a (ATCC 30-2007™) for SKBR3, RPMI-1640 (ATCC 30-2001™) for HCC1143 and DMEM (ATCC 30-2002™) for MCF-7 with 0.01 mg/mL insulin (Sigma-Aldrich^®^ Cat. No. I9278). Cells at 70–80% monolayer confluency were washed with Versene (Thermo Fisher Scientific Cat. No. 15040-066), detached with Trypsin-EDTA 0.25% (Thermo Fisher Scientific Cat. No. 25200-056), collected and counted. For each cell line, chamber slides (BioCoat™ Collagen I 8-Well Culture Slide, Corning^®^ Cat. No. 354630) were used to seed cells at a concentration of 0.1e6/mL in 0.5 mL per chamber and grown in 37 °C, 5% CO2, 100% humidity for 48 hours prior to drug compound addition. Each breast cancer cell line was seeded using individual chamber slides to avoid any cell cross-contamination, and each IMC experiment was carried out independently. In chronological order, seeding of HCC1143 was the first sample preparation, then we run the second experiment using SKBR3, and last we used MCF-7 for the drug-treated cells in vitro study.

### *In vitro* drug treatment

All chemical compounds were dissolved and aliquoted at the supplier’s recommended concentrations for long-term storage in 100% DMSO (Sigma-Aldrich Cat. No. 276855). Final concentrations for each compound were selected from previously published data [32, 53, 54] by premixing the required initial drug stock volume with full growth media, then transferring 0.5 mL volume to corresponding cell chambers (**see Supplementary Table S1**): 10 μM of etoposide (Cell Signaling Technology® Cat. No. 2200S); 0.5 μM of nocodazole (Sigma-Aldrich Cat. No. M1404); 10 ng/mL of human EGF (Sigma-Aldrich Cat. No. E9644); 2% DMSO in growth media. One chamber was left with fresh growth media as non-treated control. Cells were exposed to compounds for 48 hours at 37 °C, 5% CO2 and 100% humidity. Preparation of additional eight chamber sample slides to replicate our experiment for IMC analysis was not required due to the known *in vitro* predicted effect of the drugs we tested on this cell line.

### Immunocytochemistry and antibody validation

Immunostaining of live cells was performed in chamber slides. Following drug treatment, fresh media in all chambers was supplemented with Cell-ID™ Intercalator-103Rh (Fluidigm Cat. No. 201103A) at 1:500 for 15 minutes at 37 °C, 5% CO2 for dead cell identification. The chamber slides were washed with DPBS (Thermo Fisher Scientific Cat. No. 14190-144) at room temperature (RT) for 5 minutes. Surface metal-labeled antibody cocktail (**see Supplementary Table S2**) was made in 0.5% BSA/DPBS from antibody stocks at a dilution of 1:100 for each. Cell monolayers were stained with 250 μL antibody mix for 120 minutes, RT. Following a washing step, cells were fixed with 250 μL freshly made 1.6% formaldehyde (Thermo Fisher Scientific Cat. No. 28906) in DPBS for 15 minutes at RT. After fixative removal, cells were permeabilized with fresh 0.3% Saponin/DPBS (Sigma-Aldrich Cat. No. S7900-25G) for 30 minutes at RT, then blocked with 1% BSA/DPBS for 1 hour at 37 °C. After blocking, cells were incubated overnight at 4 °C in 250 μL freshly prepared cocktail of metal-labeled antibodies targeting intracellular markers (**see Supplementary Table S3**) diluted 1:50 in 0.3% Saponin/DBPS. The following day cells were washed once with 0.5% BSA and stained with 250 μL of nuclear Cell-ID Intercalator-Ir for 30 minutes (1:1000), RT. After the last wash, each chamber was filled with 250 μL deionized water for 5 minutes for salt removal. Chambers were detached from slides and left to dry at RT until IMC analysis. The commercial antibodies of our panels have been previously validated by immunocytochemistry applied to immunofluorescence analysis of adherent cells. A similar sample preparation method was applied to the validation of these antibodies labeled with metals regarding their antigen specificity detected by IMC. The selection of titers tested for surface markers (1:100) and intracellular markers (1:50) refer to rigorous Quality Control antibody validation functional assays using single-cell suspension Mass Cytometry, which has the same single isotopic detection modalities of antigens as IMC. The list of antibodies used for each breast cancer cell line experiment is cross-referenced in Supplementary Tables S2 and S3. A total number of 14 antibodies were tested on HCC1143 cells, 20 markers for SKBR3 and 25 for MCF-7. The markers from each panel were selected based on their specificity to surface membrane, cytoplasmic and nuclear components of single cells, their previous applications in the context of breast cancer cell line studies, and their commercial availability. Expanding a panel to 37 markers requires additional custom labeling of target specific monoclonal antibodies with non-used metal isotopes.

### Data acquisition by IMC

Samples were analyzed with the Hyperion Imaging System (Fluidigm). The dried slide was loaded into the imaging module, where an optical previewing of the ROIs was recorded for laser ablation. Areas of dimension 1000 × 1000 μm were acquired for HCC1143 and 1400 × 1400 μm for SKBR3 cell lines experiments. Replicate ROIs of 1500 × 1500 μm size were collected for MCF-7 exposed to chemical compounds, and a single ROI was ablated for non-treated control and DMSO-treated cells. Each ROI sample acquisition took 4.5 hours at an ablation frequency of 200Hz. The resulting data files were stored in MCD binary format. Multicolor images were generated with open source ImageJ 1.51 software [55]. Zoom-in regions for each multicolor image were made by selecting and cropping areas of dimension 400 × 400 μm from the original picture without any change of the pixel size. The resolution of each IMC image shown is 1 μm for 1 pixel.

### Image analysis software tools

For each recorded ROI, stacks of 16-bit and 32-bit single-channel TIFF files were exported from MCD binary files using MCD™ Viewer 1.0 (Fluidigm). Cell-based morphological segmentation was carried out with two image processing pipelines for 16-bit or 32-bit TIFF files and CellProfiler 2.2.0 (CP), a widely adopted software in the open source image analysis community which has been continually improved since its availability in 2005 to read and analyze cell images using advanced algorithms [56]. The 16-bit TIFF unstacked image format was used as input data in CellProfiler to run a segmentation pipeline with a set of sequential modules to generate and save an unsigned 16-bit integer single cell mask TIFF image. Inputs of 16-bit TIFF images with their corresponding segmentation mask were uploaded in histoCAT to open a session data analysis. Due to the high number of 29 single isotopic channel TIFF images for each replicate and drug treatment condition, histoCAT analysis was generated faster with a 16-bit format instead of a 32-bit image format which is bigger in size. The 32-bit TIFF unstacked images were loaded in CellProfiler to run a segmentation and mean intensity multiparametric measurement analysis of individual cells. The resulting outputs were exported and saved as SQLite single-cell object measurements database files directly compatible with CellProfiler Analyst for downstream computational analysis [37]. Spatial distribution maps, dimensionality reduction and unsupervised clustering for 16-bit single images were performed using the histoCAT 1.73 open source toolbox [21]. Supervised analysis by support vector machine learning classification on 32-bit datasets was processed with CellProfiler Analyst 2.2.1 [38]. Hierarchical clustering and correlation heat maps of classified cell populations were prepared using the web-based tool Morpheus (GenePattern, Broad Institute) [39]. Clustering and visualization of different types of markers in classified populations of cells across all controls and drug treatments were performed with the free software environment R 3.5.3, the packages pheatmap 1.0.12, igraph 1.2.4 and edgebundleR 0.1.4 [58].

### Statistical analysis

Data from non-treated control, DMSO-only and drug-treated cells were analyzed using a non-parametric Mann-Whitney test with a two-tailed P value for every channel independently. Statistical significance was defined as P <0.05. All statistical analysis was performed with GraphPad™ Prism^®^ 7.04 software [57].

## Results

### Multiparametric characterization of breast cancer cell lines by IMC

IMC analysis revealed heterogeneity in the expression of surface, intracellular and nuclear markers in human breast cancer cell lines cultured in chamber slides. As an example, we selected the SKBR3 cell line, which is characterized by an invasive phenotype and increased proliferation *in vitro* and is used as a model to delineate mechanisms of resistance to ErbB2-targeted clinical therapies [24, 25]. As seen in **Fig. 2**, the culture consists of mostly small and medium-size rounded epithelial cells and occasional large multinucleated cells. Most cells show high levels of the cytoskeletal marker pan-keratin. Human epidermal growth factor receptor 2 (HER2) and EGFR are seen at the surface membrane, while tumor suppressor and transcription factor p53 is localized in the nuclei. Presence of lysosomal organelles is outlined in the cytoplasm with an antibody targeting the lysosome-associated membrane protein 2 (LAMP-2), known as CD107b. Image resolution is similar to fluorescence microscopy. Ductal breast carcinoma-derived triple-negative HCC1143 cell line was used to demonstrate mesenchymal morphology (**Fig. 3**) with high vimentin detection in the cytoplasm, nuclear localization of p53 and cytoplasmic distribution of the basal-like breast cancer marker cytokeratin 5 [26, 27]. The proliferation marker Ki-67 identifies cells in the active state of the cell cycle. Combination of multiple markers in an *in vitro* cell screening assay allows more precise analysis of cellular states of differentiation and invasiveness characteristic for breast cancer tumors.

**Figure 2.**
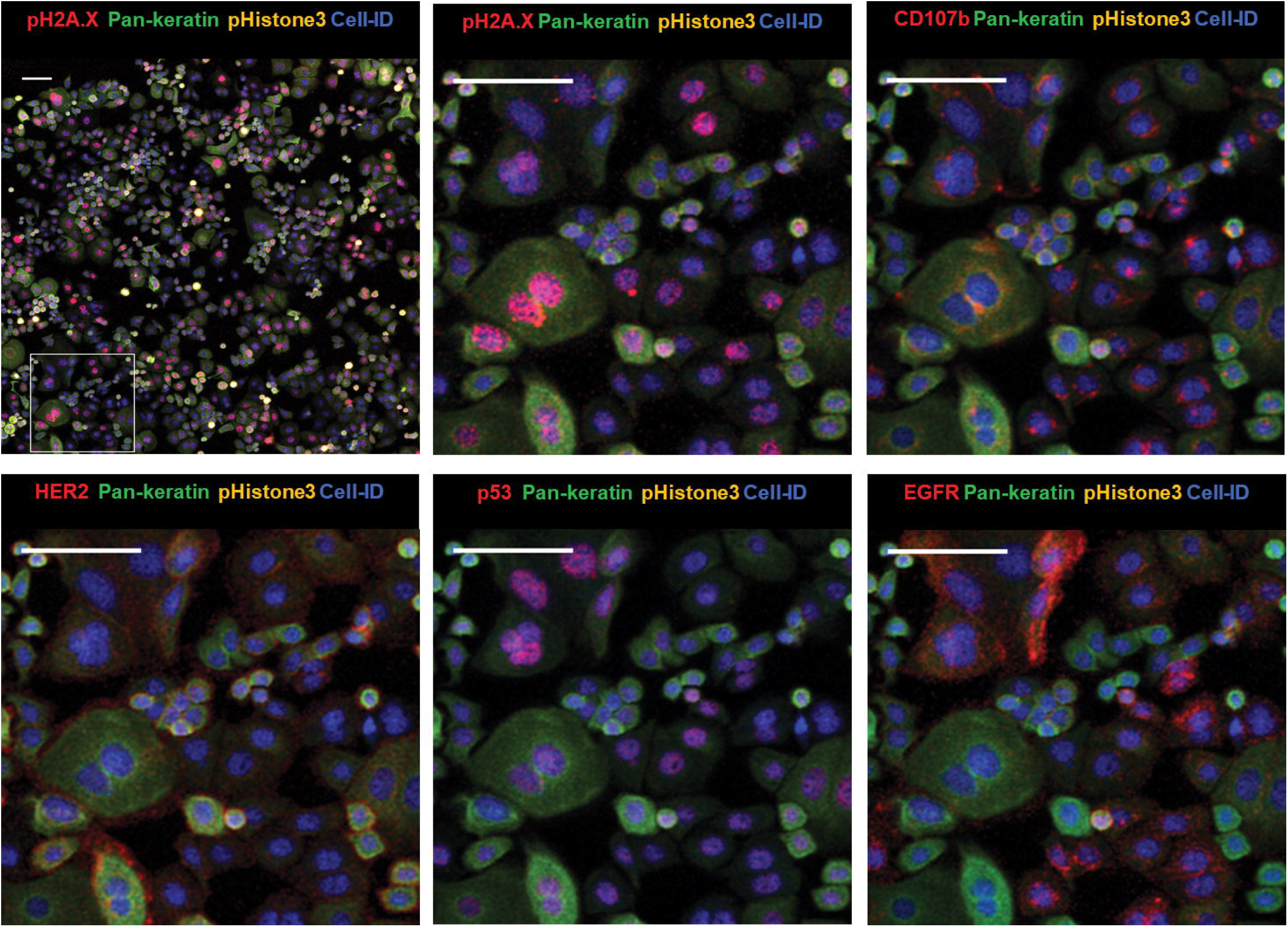
IMC images of cellular and nuclear markers expressed by SKBR3 with zoom-in area. Cell-ID: nuclei labeled with Ir-intercalator. Zoom-in images are rendered with multiple combinations of protein markers detected in different cellular compartments. Scale bar = 100 μm.

**Figure 3.**
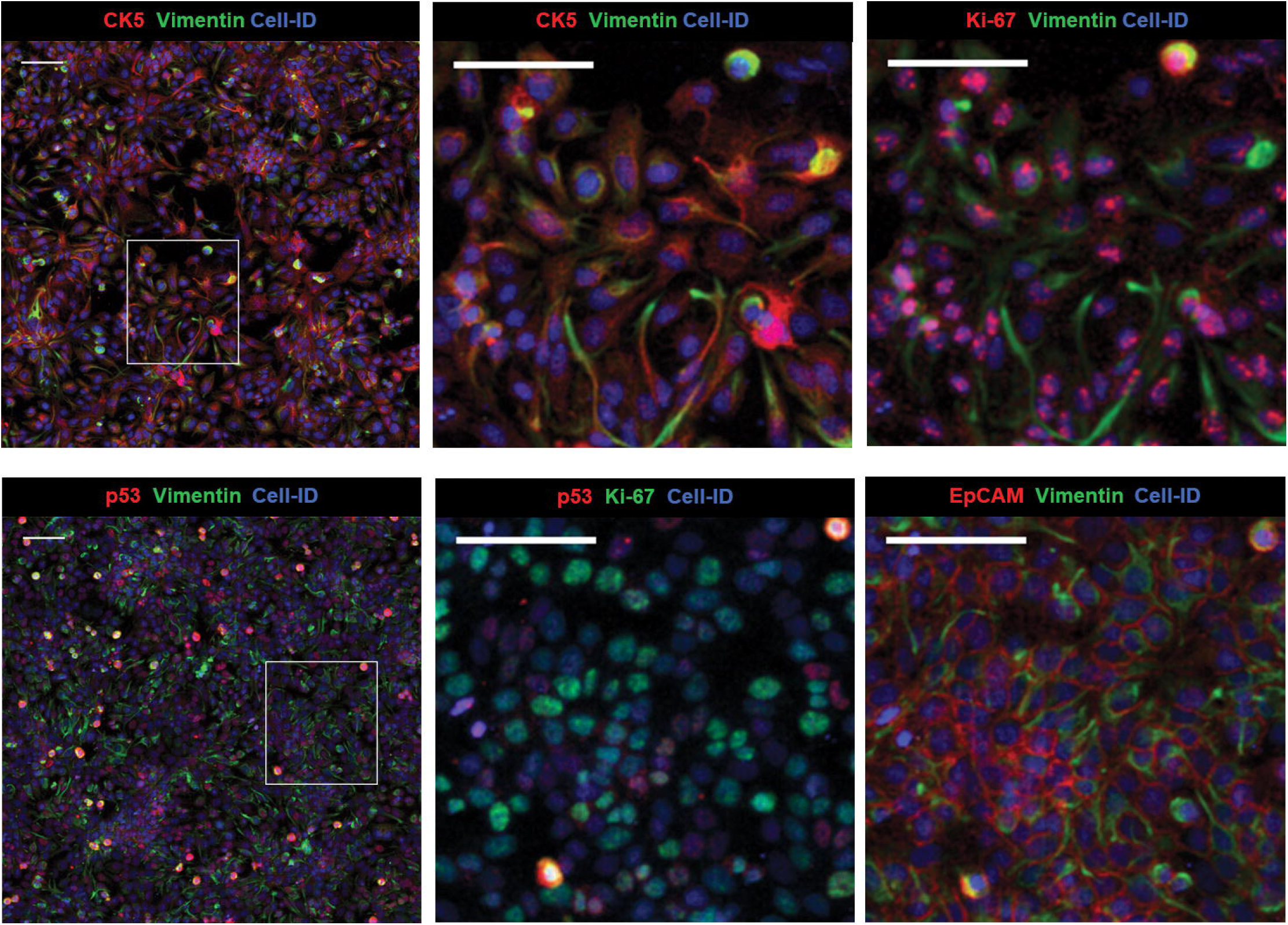
IMC images of cellular and nuclear markers expressed by HCC1143 with zoom-in areas (two ROI). Cell-ID: nuclei labeled with Ir-intercalator. Each zoom-in composite image is rendered with a selection of different markers. Scale bar = 100 μm.

### *In vitro* drug effect profiling by IMC

To evaluate how IMC may be translated into future applications in the field of cell-based drug screening, we initiated a limited drug treatment study using one model breast cancer cell line. The phenotypic and functional responses of wild-type p53-expressing MCF-7 cells to etoposide, nocodazole or EGF compounds were investigated. In preclinical *in vitro* drug screening, this model is considered to be reproducible, fast and inexpensive despite the lack of clinical correlation to drug response *in vivo* due to breast tumor heterogeneity [28]. To identify how these chemical compounds induce cytostatic and cytotoxic effects on this noninvasive luminal breast cancer cell line, we generated multiplexed pseudocolor images of affected cellular compartments (**Fig. 4**). For example, nuclear size and proliferative activity of each cell were analyzed using the nuclear intercalator-Ir stain (Cell-ID Intercalator-Ir), cell cycle-S phase marker (Cell-ID IdU), proliferation marker Ki-67, cell cycle regulatory proteins cyclin B1 and D3 and estrogen receptor alpha (ERα). Antibody against pH2A.X^S139^ was used as a nuclear marker for DNA double-strand breaks and pHistone3^S28^ for mitotic cells. Cytoskeletal markers pan-keratin and cytokeratin 19 were used to follow changes in morphology and cell size. Drug response of MCF-7 morphology, adhesion and cell-to-cell interaction were visualized with surface membrane proteins, such as the membrane-bound mucin marker MUC-1 overexpressed in breast carcinoma [29], cell adhesion molecule EpCAM, integrin CD29, tetraspanin CD81, the low-expressed epidermal growth factor receptor EGFR, regulators of integrin-mediated cell adhesion CD98 and CD47 [30], and E-cadherin as epithelial interaction protein (**Supplementary Table S2**).

**Figure 4.**
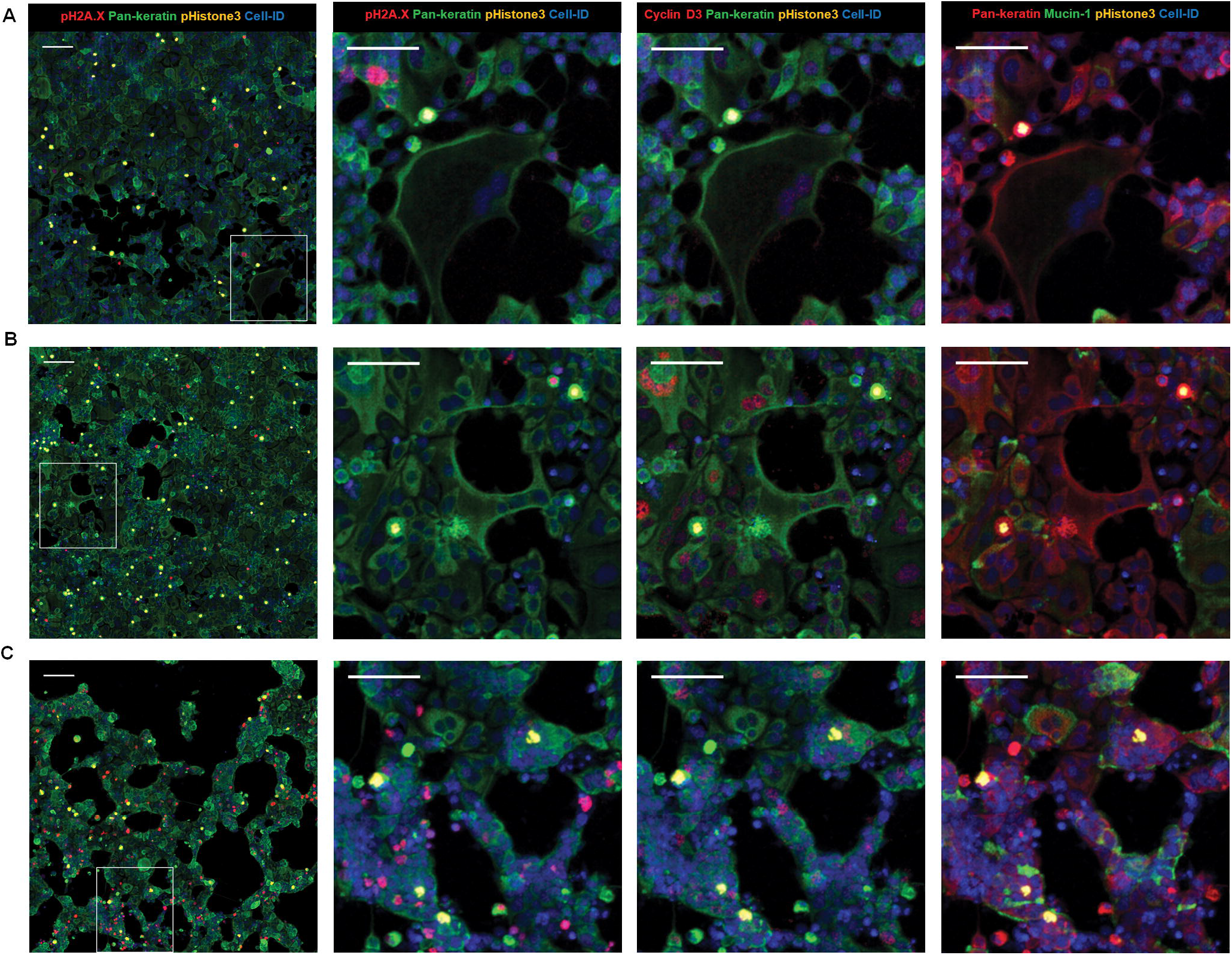
Multiplexed IMC images of MCF-7 cells treated with different compounds. EGF, (B) nocodazole, (C) etoposide. Composite images of 1500 × 1500 μm size (scale bar = 160 μm) with 400 × 400 μm zoom-in areas (scale bar = 100 μm). Columns images correspond to the same combination of proteins detected per drug treatment (row).

Long-term exposure of MCF-7 cells to EGF induced significant morphological changes and an increase in cell size. Lineage shift from a classic epithelial cobblestone monolayer to a more invasive mesenchymal type was observed. The presence of large spindle-shaped cells that seemed to lose contact with other cells and migrate into empty spaces may be related to an increase in cell motility (**Fig. 4A**). Activation of the EGFR signal transduction pathway by EGF is known to play a role in cellular motility and size increase in the tumor aggressiveness [31]. Screening of MCF-7 cells treated with etoposide, nocodazole or EGF shows multivariate phenotypic response profiles compared to non-treated and DMSO-treated controls (**Supplementary Fig. S1**).

Treatment of MCF-7 cells with the antineoplastic agent nocodazole resulted in an increase in the population of pHistone3^S28^-positive cells with a mitotic index three-fold higher compared to DMSO-treated or non-treated control (**Supplementary Table S4**). Another cytostatic phenotype was observed after prolonged drug exposure: subpopulations of cyclin D3-positive and tetraploid cells exiting mitosis without cytokinesis as a result of microtubule disruption by nocodazole (**Fig. 4B**). Presence of cyclin D3-positive and tetraploid cells in the G1/S interphase confirms nocodazole effect on MCF-7 cell cycle [32].

Incubation of MCF-7 with the anti-cancer agent etoposide shows inhibition of cell proliferation, elevated numbers of cells positive for nuclear pH2A.X^S139^ protein and compaction of the monolayer compared to EGF- or nocodazole-treated samples (**Fig. 4C**). Exposure of tumor-derived cell lines to this inhibitor induces the formation of a stable covalent topoisomerase II-cleaved DNA complex, which causes multiple breaks in double-stranded DNA. Etoposide treatment of MCF-7 activates phosphorylation of H2A.X, which is required for checkpoint-mediated cell cycle arrest, promotes DNA repair and maintains genomic stability. Thus, pH2A.X^S139^ is a sensitive biomarker for measuring levels of drug-induced double-stranded DNA breaks in cancer cells [33].

### High-dimensional phenotype analysis of *in vitro* drug response

Multivariate phenotypic responses of MCF-7 cells following chemical perturbations were quantified with a set of available software tools. Unfortunately, there is no single software package for this type of analysis. Therefore, we employed a workflow that includes a wide variety of tools designed for cellular imaging. Hopefully, this work will underscore the need for software specifically developed for analysis of multiparametric data acquired by IMC.

First, we used CellProfiler 2.2.0 (CP) to identify single cells through establishing nuclear and cellular boundaries [34]. Accuracy and robustness of cell segmentation was assessed by using Cell-ID Intercalator-Ir to identify cell nuclei and pan-keratin for cytoplasm (**Fig. 5A, Supplementary Fig. S2A and S3**). The segmentation masks were generated for each ROI, then exported for further downstream analysis of changes in MCF-7 cellular phenotypes in response to drug treatment.

**Figure 5.**
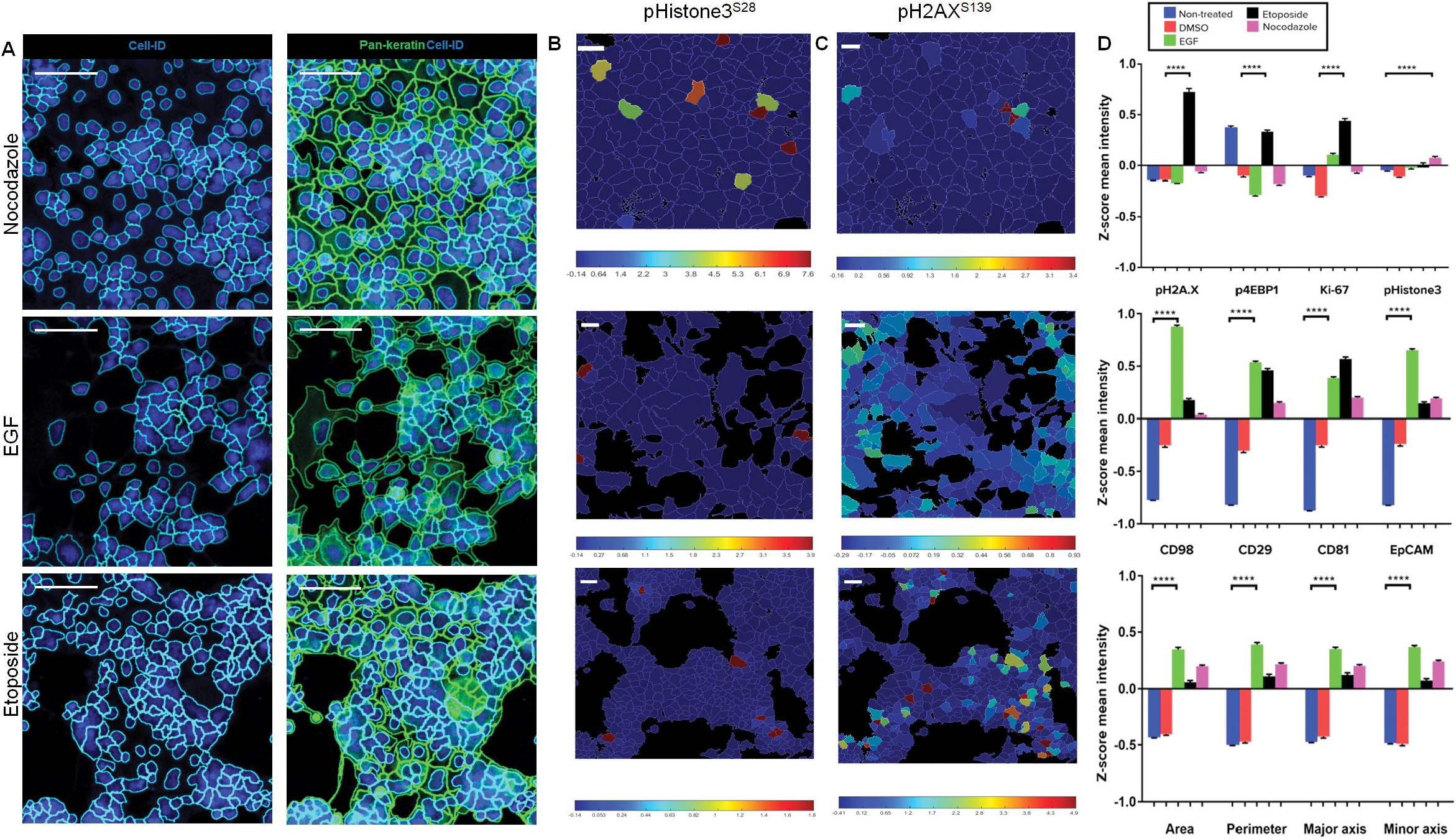
Image processing by cell-based segmentation and spatial parametric distribution across ROIs of drug-treated MCF-7 cells. (A) Cell and nuclear masks overlaid on color IMC images. Scale bar = 100 μm. Zoom-in areas of selected parameters visualized by Z-score normalized color heat maps for (B) pHistone3^S28^ (C) pH2A.X^S139^. Scale bar = 10 μm. (D) Z-score mean intensity levels of nuclear markers (top), surface markers (middle) and cellular size parameters (bottom) for control and drug treatment conditions. Data are presented as mean ± SEM. ****P < 0.0001 (unpaired t-test).

Second, to visualize and analyze data we used the open source platform histoCAT, which enables highly multiplexed, quantitative and detailed analysis of cell phenotypes, microenvironment interactions and tissue architecture [21]. The workflow of histoCAT consists of overlaying the segmentation masks previously generated by CP to extract single-cell level information.

To evaluate the diversity of phenotypic profiles collected from drug-treated MCF-7 cells, we then selected and generated Z-score normalized mean intensities of pHistone3^S28^ and pH2AX^S139^ as drug-sensitive protein markers. Spatial heat maps of both channels were displayed for various drug-treated MCF-7 ROIs using a 99^th^ percentile cut-off (**Figs 5B and 5C, Supplementary Fig. S2B**). This quantitative and visual approach allows us to compare at the same scale the spatial distribution of these parameters between ROIs. For example, pH2A.X^S139^ nuclear expression level in etoposide-treated MCF-7 cells is higher than in nocodazole or EGF-treated cells on average. Mitotic cell marker pHistone3^S28^ expression level in nocodazole-treated MCF-7 is two-fold higher than those identified for other drugs. Related single-cell Z-score mean intensity values for nuclear markers (pHistone3^S28^, pH2A.X^S139^, Ki-67, p4EBP1^T37/T46^, p53, cyclin D3), surface markers (CD98, CD81, CD29, CD49e, CD47) and cell size parameters (area, perimeter, major and minor axis lengths) were exported from each drug treatment dataset to assess statistically significant changes of protein expression levels and cellular morphology (**Fig. 5D and Supplementary Fig. S4)**. We can see differences between both controls and drug-treated samples in the expression of nuclear markers, with a co-increase of pH2A.X^S139^ and Ki-67 levels, and phosphorylation of the translation repression protein 4E-BP1 under etoposide treatment. This suggests the presence of a non-quiescent population of MCF-7 undergoing DNA repair and maintaining high proliferative activity. For nocodazole and EGF treatments, low Ki-67 levels may be related to the disruption of cell cycle and lineage shift induced in response to respective compound [35]. EGF treatment shows increased cell size and expression of adhesion surface markers compared to other conditions. These quantitative results are in agreement with multicolor images of treated cells.

Finally, the heterogeneity of MCF-7 cell phenotypes was analyzed using the t-distributed stochastic neighbor embedding (t-SNE) algorithm, a data dimensionality reduction method based on the similarity of selected markers or cell types [36]. We processed replicates of ROI for each compound using Z-score-normalized protein markers and cell size parameters to generate two-dimensional t-SNE plots. This visualization tool shows signal distribution of phosphoproteins, size parameters and nuclear and cell surface membrane markers over the different ROIs (**Fig. 6A and Supplementary Fig. S5**). For example, the population of mitotic cells was grouped in a single region on the map with high expression levels of p4EBP1^T37/T46^, pHistone3^S28^ and Ki-67. The high level of pH2A.X^S139^ was detected in the region of etoposide-treated cells and correlates with the strong Ir-intercalator signal due to compaction of cellular DNA in response to the drug. Area plot reveals the presence of larger-size cells in EGF and nocodazole treatments, in line with EGF-induced lineage shift and cytokinesis inhibition by nocodazole (**Fig. 6B**).

**Figure 6.**
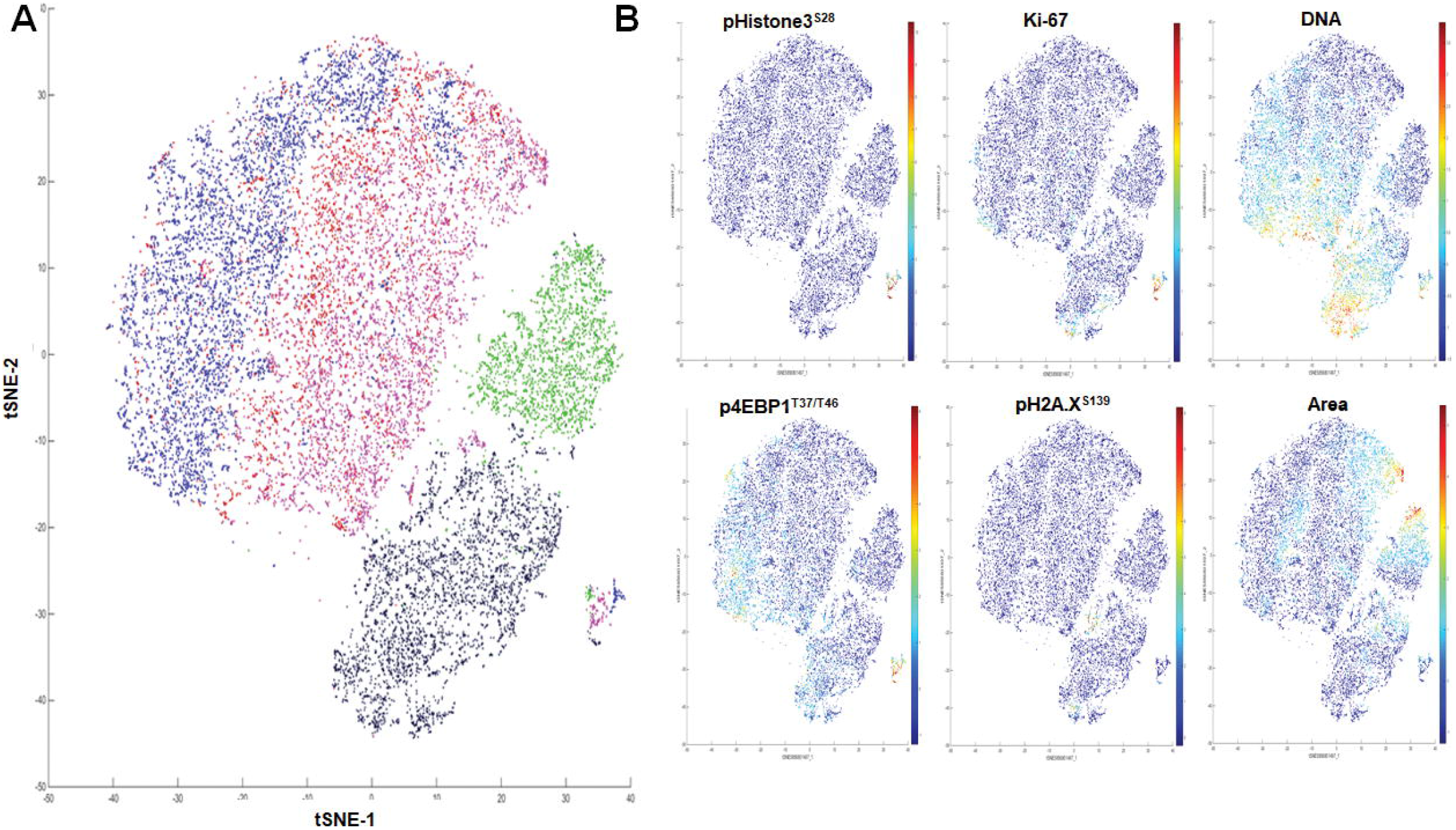
Unsupervised cluster identification by t-SNE of control and compound-treated MCF-7. (A) Combined colored t-SNE maps of individual cells (EGF, green; etoposide, black; nocodazole, purple), DMSO (red) and non-treated control (blue). (B) t-SNE maps colored according to the expression level of nuclear markers and cell size (area).

### Classification of drug-treated MCF-7 phenotypic changes by support vector machine learning

Here we applied the support vector machine learning classification ruler termed Fast Gentle Boosting for accurate IMC dataset multiparametric analysis of nuclear states of compound-treated MCF-7 cells [37]. This classifier was integrated into CellProfiler Analyst (CPA) and trained [38] to identify five different nuclear classes of MCF-7 cells: pHistone3^S28^-positive mitotic cells, pH2A.X^S139^ for cells undergoing DNA repair, p53 as tumor suppressor activation, cyclin D3 for G1/S transition in interphase and Ki-67 for proliferating cells. A composite image of all nuclear markers is shown in **Fig. 7A**. These single-cellular objects were then scored and classified for a supervised analysis of every protein marker and cell size parameter per specified nuclear subpopulation. Accuracy of image classification on each dataset of MCF-7 drug compound was evaluated by cross-validation between the true labeled cells as classification training set and the image full scoring as predicted label set for each nuclear class. Classification reports were displayed as confusion matrices where each value corresponds to average cross-validation metrics between the trained (true label) and scored (predicted label) single-cell objects for each nuclear class on a performance scale from 0 to 1 (**Fig. 7B and Supplementary Fig. S6**). As shown, classification of each nuclear class per drug compound dataset shows relatively good performance between 0.5 and 1.

**Figure 7.**
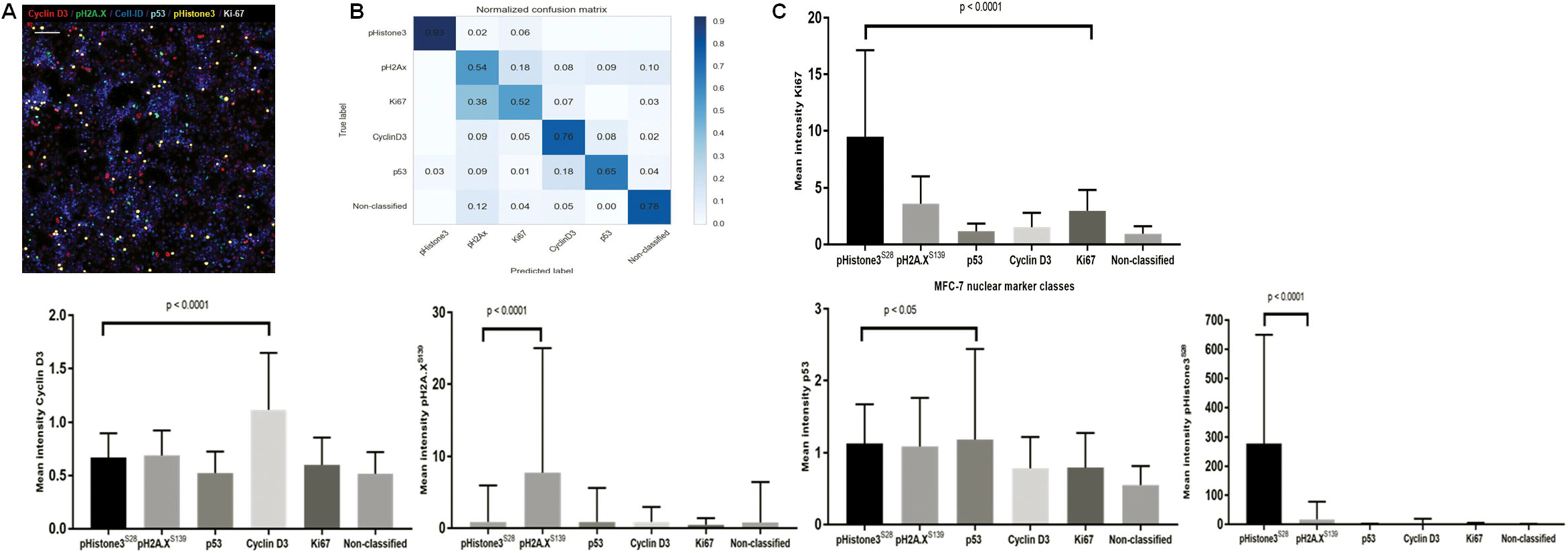
Support vector machine learning classification and multiparametric hierarchical similarity clustering of drug-treated MCF-7. (A) 6-plex color image of nocodazole-treated cells generated by CPA Image Viewer. Scale bar = 160 μm. (B) Normalized confusion matrix of six different cell nuclear phenotypes. (C) Protein expression levels of nuclear markers used for cell classification of MCF-7 treated with nocodazole. Graphs represent the non-normalized mean intensity (±SD) of nuclear proteins within each assigned category. P < 0.05 and P < 0.0001 (unpaired t-test) are included.

Raw mean intensity values of protein expression for each of the five nuclear markers were exported per classified MCF-7 population and compared to evaluate statistical significance (**Fig. 7C and Supplementary Fig. S7**). Bar charts show various levels of protein markers among the five predicted classes of nuclear states, with higher protein detection in the corresponding class, except for the Ki-67 proliferative marker, which shows stronger detection in pHistone3^S28^ mitotic cells.

Multiparametric MCF-7 drug compound data pre-labeled by nuclear classification were reprocessed using hierarchical clustering similarity heat maps. First, we computed pairwise similarities for each class of cells per drug exposure with Pearson correlation coefficient between each measured parameter. This method is applicable to non-normalized data and can be used when data measurements vary between samples. Similarity Pearson correlation heat maps were computed using the browser-based tool Morpheus [39]. The core interface in Morpheus is a graphical color heat map representing protein marker and size parameter measurements applied to each single-cell object. Determination of the Pearson correlation coefficient between each measured parameter generates a reorganized heat map of multiparametric similarity relationships for each classified MCF-7 drug dataset [40]. Using these hierarchical similarity clustering heat maps, we compared the grouping pattern of mitotic pHistone3^S28^-positive cells across the multiple ROIs of compound-treated MCF-7 (**Fig. 8 and Supplementary Fig. S8**). Naturally occurring isotopes of iridium, ^191^Ir and ^193^Ir, are present in the nuclear staining reagent Cell-ID Intercalator-Ir and are grouped consistently across all heat maps. The two cytoskeleton biomarkers pan-keratin and cytokeratin 19 strongly correlate as well, since anti-pan-keratin and anti-CK19 antibodies identify cytokeratin isoforms highly expressed in MCF-7. Size parameters including perimeter, area and axis length of each mitotic cell are grouped together after hierarchical clustering across all drug treatments.

**Figure 8.**
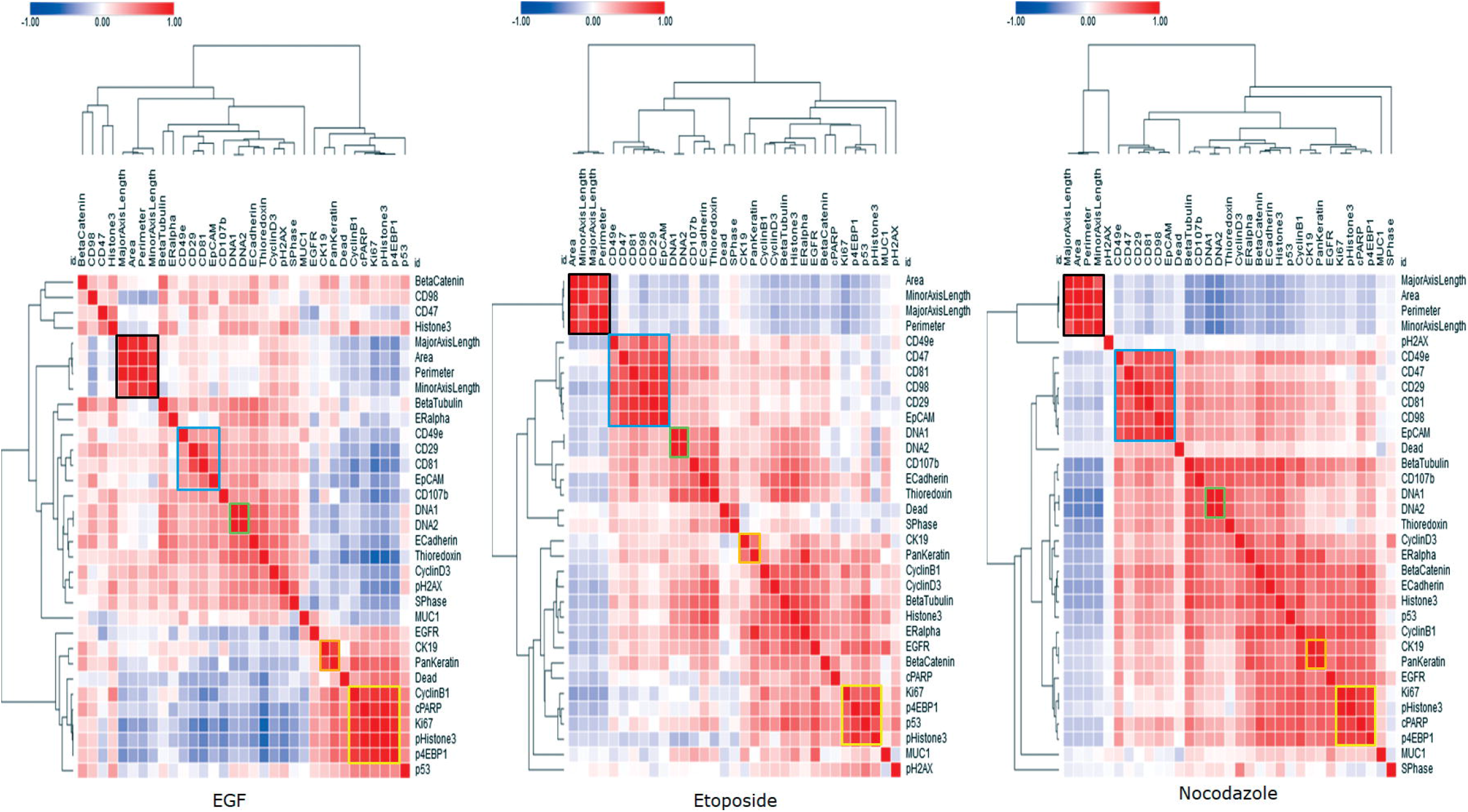
Hierarchical clustering heat maps of Pearson coefficients of multiple parameters for drug-treated pHistone3^S28^-positive cells. Clusters of cells are highlighted according to cells size parameters (black), CD surface markers (blue), Cell-ID Intercalator-Ir isotopes (green), pan-keratin and CK19 markers (orange) and nuclear markers (yellow).

Cell surface protein markers (CD29, CD98, CD81, CD47, EpCAM and CD49e) generated a cluster with high correlation and redundancy in the pHistone3^S28^ mitotic cell subclass. However, EGF treatment influenced expression of these markers, and some level of dissimilarity occurred between the surface membrane cluster and two integrin-mediated regulators, CD98 and CD47. This is consistent with the fact that MCF-7 cells switch to a more mesenchymal phenotype after EGF treatment and higher motility, with a concomitant increase in the expression level of some adhesion proteins.

The intracellular and nuclear biomarkers pHistone3^S28^, p4EBP1^T37/T46^ and Ki-67 for mitotic cells cluster together in all conditions, showing a specific phenotypic profile. By looking at the correlation between the mitotic marker pHistone3^S28^ and other markers, we see that p53, cyclin B1 and cleaved poly (ADP-ribose) (cPARP) show variations in their correlation similarities depending on the drug treatment. p53 and cPARP exhibit close relationships with other nuclear markers, since both bind to nuclear DNA to be functional. Cyclin B1 shows strong correlation with pHistone3^S28^ in cells from non-treated control and EGF-treated ROIs. This correlation is less pronounced in the case of etoposide and nocodazole.

The clustering of different types of markers in the class of pHistone3^S28^ mitotic cells was measured by comparing the similarity level of each target-specific cluster across all controls and drug treatments. First, we reprocessed cell surface membrane CD markers for each treatment as a new hierarchical similarity heatmap by using Pearson correlation as distance metric and complete linkage method (**Fig. 9A**). As expected, each individual cluster is aligned and identified around the matrix diagonal component. Both non-treated and DMSO controls group of protein markers show close similarities to each other, and low or negative correlation with drug-treated markers. We used a hierarchical visualization edge bundle circle plot diagram to highlight the positive intra and inter-similarities across surface membrane protein parameters (**Fig. 9B**). Visualization of the high pairwise correlations within each cluster is shown as edge connecting nodes, and bundled lines correspond to close inter-relationships between both control conditions. Drug-treated clusters of proteins show poor or no interdependency. A similar visual workflow was applied to the multivariate analysis by pairwise Pearson correlation of specific nuclear markers and cytoplasmic markers types (**Supplementary Fig. S9**).

**Figure 9.**
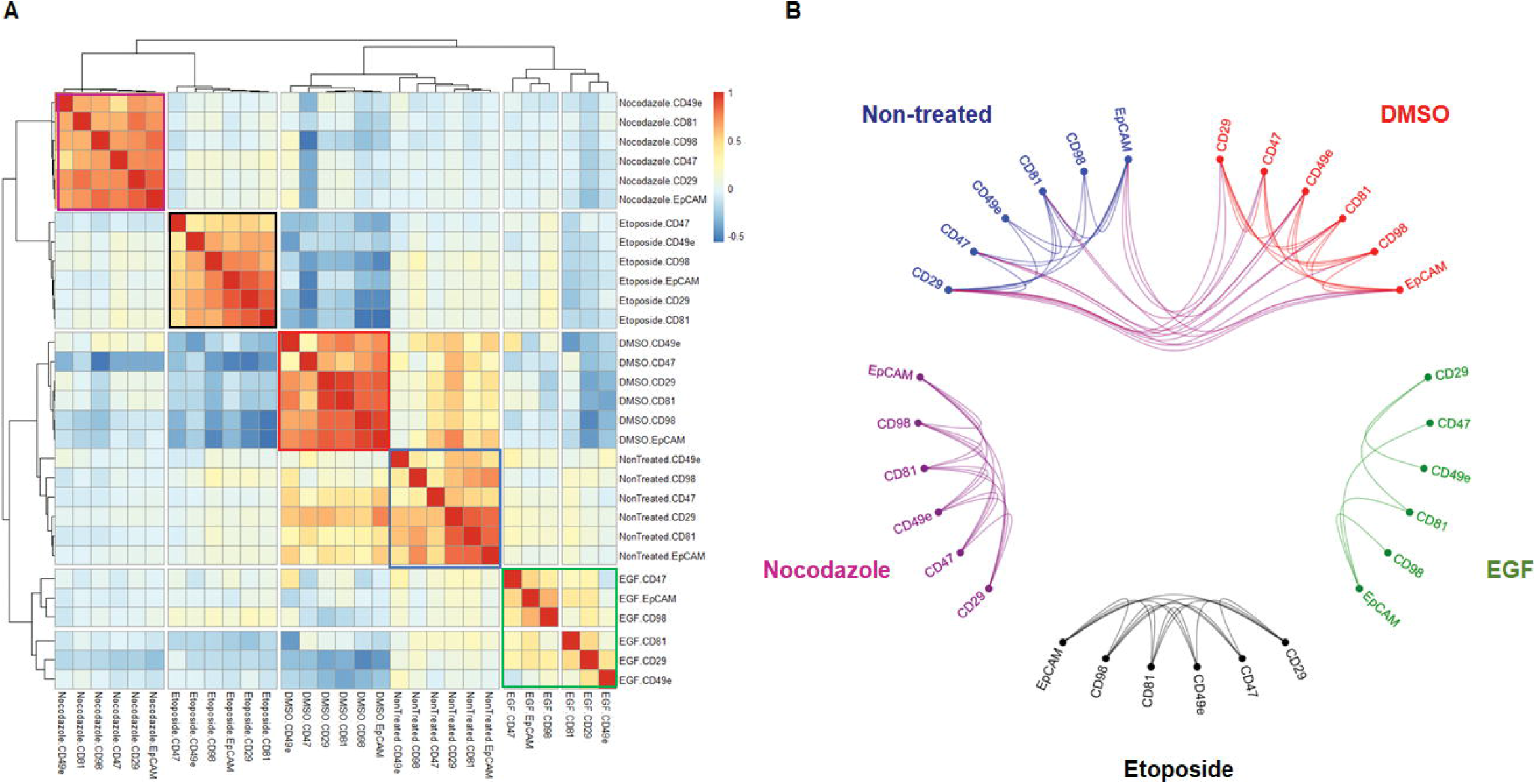
Comparative analysis of multiparametric pairwise correlations between surface membrane markers expressed by MCF-7 pHistone3^S28^ positive mitotic cells across controls and drug treatments. (A) Hierarchical similarity heatmap of CD98, CD29, EpCAM, CD81, CD47 and CD49e protein markers. Individual clusters are highlighted around the diagonal component for each condition (Non-treated, blue; DMSO, red; EGF, green; etoposide, black; nocodazole, purple). (B) Hierarchical edge bundle visual graph showing connecting edges between parametric nodes. Bundled lines between nodes correspond to positive pairwise correlations with a cut-off value > 0.3. The bundled interconnections show similarities of surface membrane phenotyping profiles between non-treated and DMSO controls.

## Discussion

The goal of this study was to demonstrate a multidimensional IMC workflow and quantitative data processing using open source computational tools for *in vitro* drug response research. The multiplexed measurements acquired from a single image may provide a more informative and reliable screening of drug leads for further clinical development [41]. In this work, phenotypic features of tumor-derived cell lines such as SKBR3, HCC1143 and MCF-7, which recapitulate different subtypes found in breast tumors, were characterized. The wild-type p53-expressing MCF-7 epithelial cell line, a model of luminal breast adenocarcinoma, was selected for *in vitro* drug treatment and exposed for 48 hours to three classes of bioactive compounds: EGF, nocodazole and etoposide. The cells were stained with a mixture of metal-tagged antibodies against cell membrane, cytoplasmic and nuclei markers and subjected to IMC. The panel of metal tag antibodies used in our study was designed to minimize any potential spill over cross-interference between neighbor isotopic mass channels [42, 43]. When analyzing our IMC data, we did not encounter strong and significant channel contamination from one metal labeled antibody to another. The collected data allowed us to develop a workflow for cellular imaging analysis, which relies on established computational methods [44]. Multiplexed images of cell compartments give a detailed and broad visualization of biomarker localization and cell size parameter changes. Cell-based morphological segmentation of generated digital images is a prerequisite to identify and export multiple features of interest at single-cell resolution. Exploratory downstream analysis was performed with the histoCAT platform to generate, normalize, measure and compare multiple images according to their multidimensional content in the form of spatial distribution heat maps, dimensionality reduction and unsupervised clustering. In this report, we also describe cell classification by machine learning to pre-label cells before measuring their multivariate similarities. Prior knowledge of different cellular states is important for the calculation of distance and similarity metrics between each marker in the evaluation of the heterogeneity of cells and the effects of different compounds on cancer cells [45]. In the example of pHistone3^S28^-positive mitotic cells, we generated a multivariate high-dimensional heat map that allows a comparative study of mechanisms of action of multiple drugs. Additional nuclear classes such as pH2A.X^S139^, p53, cyclin D3 and Ki-67 were analyzed using similar multiparametric clustering approaches (data not shown). The strategy could enable new ways to perform deep proteomic drug screening on different models for translational medicine and preclinical trials [46, 47].

Other *in vitro* assays using stem cells, or co-cultures of different cell types, could be studied by IMC as an additional approach to designing high-content analysis to investigate cell-to-cell interactions and signaling. A recent targeted therapy against cancer oncogenes described a cytotoxic small-molecule compound that activated the steroid receptor signaling pathways and led to cell stress and death [48]. IMC may assist researchers to better understand perturbations in this and other pathways through high-dimensional analysis. Three-dimensional tumor spheroids grown *in vitro* are more representative of the complexity of cancer tissue than two-dimensional cell cultures and can be analyzed by IMC through the preparation of sections of FFPE pelleted spheroids [49]. We summarized in **Table 1** several advantages and challenges during preparation on slide, staining procedure, acquisition by IMC and software data analysis for three different models of sample. IMC was useful in the biodistribution study of the platinum-containing anti-cancer drug cisplatin in pancreatic adenocarcinoma xenograft model and could be applied to the emerging metal-containing chemotherapies and photodynamic therapies, which are considered as alternative approaches to classic compounds [50, 51]. Therefore, the goal for IMC in preclinical screening of metallodrugs will be to efficiently and effectively identify cellular responses of *in vitro* tumor-derived cell lines for lead optimization in target-based drug discovery [52]. A comparison of IMC performance metrics to other high content imaging system technologies in the field of single cell analysis is shown in **Table 2**.

**Table 1.** Key similarities and differences of biological samples processing used for IMC.

| Sample type | Tissue sections<br>(Frozen and FFPE) | Cell smear on microscopic<br>slide (Liquid biopsy) | Adherent cell lines grown in<br>chamber slides |
| --- | --- | --- | --- |
| **Sample quality and integrity**<br>**criteria** | Thin thickness of the cut tissue slice (5µm).<br>Effectiveness of tissue fixation for FFPE (temperature, fixing chemicals, pH, time).<br>Storage conditions for tissue preservation (temperature, humidity, light exposure). | Cell monolayer smear on slide by surface tension.<br>Fixation method.<br>Low temperature storage conditions.<br>Pre-identification of cells by immunofluorescence staining and scanning. | High cell viability and proliferation rate.<br>Adherence efficiency of growing cells to the surface of the slide.<br>Optimal seeding density on slide and growth conditions per cell line tested. |
| **Sample staining critical steps** | Optimization of dewaxing and hydration procedures for FFPE tissue.<br>Removal of tissue freezing matrix for frozen tissue.<br>Antigen retrieval and blocking conditions.<br>Metal isotope labeling of validated antibody clones for tissue staining.<br>Titration range testing of metal tag antibodies and DNA metallo-intercalators. | Use of hydrophobic circle barrier around cells in specific regions for liquid staining on slide.<br>Blocking conditions.<br>Preparation of metal tag antibodies liquid cocktail.<br>Washing and drying of the sample slide before acquisition. | Metal tag antibodies panel specific to surface membrane, cytoplasm and nucleus of adherent cells.<br>Chamber well liquid volume measurements for cell staining procedure.<br>Fixation and permeabilization conditions.<br>Removal of chamber wells from the slide. |
| **Sample acquisition by IMC** | Simultaneous acquisition of all markers.<br>Whole ablation of tissue regions.<br>No autofluorescence. | Plotting of two-dimensional coordinates of cells pre-identified by immunofluorescence.<br>Laser ablation of areas with good cellular density and morphology. | Regions of interest with intact cells<br>Ablation of areas with even cell monolayers rather than densely packed overlapping cells. |
| **Data analysis challenges** | Pathology expertise required for IMC imaging data visualization.<br>Annotation of structural and cellular regions.<br>Segmentation of individual and distinct cells within tissue samples require nuclei and membrane markers. | Single-cell segmentation of small-sized cells.<br>Identification of nuclei with DNA metal-intercalator.<br>Use of membrane stain for detection of cellular boundaries.<br>Classification of cellular subpopulations. | Single-cell morphological segmentation analysis.<br>Nuclei segmentation with DNA metal-intercalator.<br>Whole cell detection using cytoplasmic marker.<br>Classification of different cell states. |
| **Multiplexing biological**<br>**applications** | Immuno-oncology<br>Solid tumor oncology<br>Quantitative pathology | Identification of circulating tumor metastatic cells<br>Hematology<br>Immunology | Drug discovery<br>Cell lines authentication and characterization<br>Single-cell systems biology |

**Table 2.** Performance comparison of IMC to other high-content single cell imaging technologies.

| High-Content Imaging Technology | Imaging Mass Cytometry | Imaging Flow Cytometry | Imaging Mass Spectrometry (SIMS MIBI, MALDI) | Single Cell Fluorescence Microscopy | Super-resolution Microscopy |
| --- | --- | --- | --- | --- | --- |
| **Reagents used** | • Metal tagged antibodies<br>• DNA metal-intercalators | • Fluorescent labeled antibodies<br>• DNA dyes | • Metal tagged antibodies (SIMS MIBI)<br>• Label free, matrix solutions (MALDI) | • Fluorescent labeled antibodies<br>• DNA dyes | • Fluorescent probes |
| **Sample format** | • Chamber slide (2, 4, 8, 16-well format)<br>• Tissue on slide<br>• Cell smear on slide | • Multi-well plates<br>• Tubes | • Tissue on slide<br>• Cell smear on slide | • Multi-well plates (96, 384, 1536-well format) | • Single coverslip<br>• Multi-well plate |
| **Sample preparation time** | 6 hours | 6 hours | 6 hours | 6 hours per staining cycle | 6 hours |
| **Spatial resolution** | 1 $\mu\text{m}$ | 1 $\mu\text{m}$ | • < 1 $\mu\text{m}$ (SIMS MIBI)<br>• 5-200 $\mu\text{m}$ (MALDI) | 1–100 $\mu\text{m}$ | < 1 $\mu\text{m}$ |
| **Throughput time** | 200 pixels per second | 1000-5000 cells per second | • 10 000 pixels per second (SIMS MIBI)<br>• 100 pixels per second (MALDI) | 100-300 cells per second | 10 cells per hour |
| **Number of cells analyzed** | $10^3$-$10^4$ | $10^4$-$10^5$ | $10^3$-$10^4$ (SIMS MIBI) | $10^3$-$10^5$ | $10$-$10^2$ |
| **Sample types** | • Fixed adherent and suspension cells<br>• FFPE and frozen tissue sections | • Cells in suspension | • Fixed adherent and suspension cells<br>• FFPE and frozen tissue sections | • Live and fixed cells | • Live and fixed cells |
| **Number of targets** | > 37 markers | 12 markers | • 7 markers per scan (SIMS MIBI)<br>• 0-100 000 Dalton m/z spectra range of molecules (MALDI) | 6 markers per scan | 4 markers per scan |
| **References** | 17, 20 | 46 | 59, 60 | 14, 60 | 60, 61 |

In conclusion, we demonstrate a comprehensive image analysis workflow for cell-based IMC data analysis of surface and intracellular markers in drug-treated cells. The correlation coefficient for pairs of multiple parameters and the similar distances among a collection of treatment profiles facilitate downstream analysis and allow for direct data visualization. Image-based cell profiling studies compute statistical estimates of the likelihood of equivalence between two drug-induced profiles. Similarity measurements quantify proximity between profiles, since they detect deviations from one sample to another regardless of the absolute magnitude. This procedure is useful in finding relations and groups of samples that share common properties in high-content screening. Predicted pharmacodynamic effects were visualized and quantified in MCF-7 cells dosed with three target-specific compounds. Strong pairwise correlation between nuclear markers pHistone3^S28^, Ki-67 and p4E-BP1^T37/T46^ in mitotic cells and anti-correlation with cell surface markers CD29, CD98, CD81, CD47 and EpCAM was demonstrated.

## Supporting information

Supplementary Informations

## Acknowledgements

The authors thank the employees of Fluidigm Canada Inc. (Markham, Ontario) for their support of this research.

## Funding

This research received no specific grant from any funding agency.

## Disclosures

A.B., A.E. and O.O. are employees of, and receive remuneration from, Fluidigm Corporation. Fluidigm, Cell-ID, CyTOF, Imaging Mass Cytometry and IMC are trademarks and/or registered trademarks of Fluidigm Corporation in the United States and/or other countries. All other trademarks are the sole property of their respective owners.

For Research Use Only. Not for use in diagnostic procedures.

## Author Contributions

AB designed the study and drafted the manuscript. AB, AE and OO took part to the manuscript writing. All authors have read and approved the manuscript.

## Supporting Information

Additional supporting information may be found online in the Supporting Information section at the end of the article.

**Figure S1.** IMC multiplex image of MCF-7 non-treated and DMSO-treated control cells.

**Figure S2.** Cell segmentation and single-marker distribution of MCF-7 non-treated and DMSO-treated controls.

**Figure S3.** Segmentation accuracy of nuclei and cytoplasmic contours of identified MCF-7 cells drug-treated using binary image overlap.

**Figure S4.** Comparison of average Z-score expression level of p53, cyclin D3, CD47, CD49e and Cell-ID Intercalator-Ir between each control and drug treatment condition.

**Figure S5.** Unsupervised cluster identification by t-SNE of control and compound-treated MCF-7.

**Figure S6.** Fast Gentle Boosting classification matrices of stitched replicates of MCF-7 ROIs per control and drug compound.

**Figure S7.** Comparative expression levels of protein markers Ki-67, cyclin D3, pH2A.X^S139^, p53 and pHistone3^S28^ (rows) per nuclear class and drug treatment condition (column) of MCF-7 cells.

**Figure S8.** Heat maps of Pearson correlation coefficients of multiple parameters for non-treated and DMSO-treated controls ROIs on mitotic pHistone3^S28^ cells.

**Figure S9.** Graphical representations of Pearson correlation coefficients for nuclei markers (A) and cytoplasmic markers (B) across all controls and drug treatments in the classified population of mitotic pHistone3^S28^ MCF-7 cells.

**Table S1.** List of chemical compounds used for MCF-7 cell treatment, 48 h exposure.

**Table S2.** Metal-labeled antibody panel for cell surface markers.

**Table S3.** Metal-labeled antibody panel for intracellular markers.

**Table S4**. Mitotic index of compound-treated MCF-7 generated by support vector machine classification with Fast Gentle Boosting ruler.

## REFERENCES

1. Masters, JR (2000) Human cancer cell lines: fact and fantasy. Nat Rev Mol Cell Biol 1:233–236.

2. An, WF and Tolliday, N (2010) Cell-based assays for high-throughput screening. Mol Biotechnol 45:180–186.

3. Wilding, JL and Bodmer, WF (2014) Cancer cell lines for drug discovery and development. Cancer Res 74:2377–2384.

4. Iorio, F, Knijnenburg, TA, Vis, DJ, Bignell, GR, Menden, MP, Schubert, M, Aben, N, Gonçalves, E, Barthorpe, S, Lightfoot, H, Cokelaer, T, Greninger P, van Dyk, E, Chang, H, de Silva, H, Heyn, H, Deng, X, Egan, RK, Liu, Q, Mironenko, T, Mitropoulos, X, Richardson, L, Wang, J, Zhang, T, Moran, S, Sayols, S, Soleimani, M, Tamborero, D, Lopez-Bigas, N, Ross-Macdonald, P, Esteller, M, Gray, NS, Haber, DA, Stratton, MR, Benes, CH, Wessels, LFA, Saez-Rodriguez, J, McDermott, U and Garnett, MJ (2016) A Landscape of Pharmacogenomic Interactions in Cancer. Cell 166:740–754.

5. Haverty, PM, Lin, E, Tan, J, Yu, Y, Lam, B, Lianoglou, S, Neve, RM, Martin, S, Settleman, J, Yauch, RL and Bourgon, R (2016) Reproducible pharmacogenomic profiling of cancer cell line panels. Nature 533:333–337.

6. Goodspeed, A, Heiser, LM, Gray, JW and Costello, JC (2016) Tumor-Derived Cell Lines as Molecular Models of Cancer Pharmacogenomics. Mol Cancer Res 14:3–13.

7. Shoemaker, RH (2006) The NCI60 human tumour cell line anticancer drug screen. Nat Rev Cancer 6:813–823.

8. Sharma, SV, Haber, DA and Settleman, J (2010) Cell line-based platforms to evaluate the therapeutic efficacy of candidate anticancer agents. Nat Rev Cancer 10:241–253.

9. Lang, P, Yeow, K, Nichols, A and Scheer, A (2006) Cellular imaging in drug discovery. Nat Rev Drug Discov 5:343–356.

10. Kang, J, Hsu, CH, Wu, Q, Liu, S, Coster, AD, Posner, BA, Altschuler, SJ and Wu, LF (2016) Improving drug discovery with high-content phenotypic screens by systematic selection of reporter cell lines. Nat Biotechnol 34:70–77.

11. Gustafsdottir, SM, Ljosa, V, Sokolnicki, KL, Anthony Wilson, J, Walpita, D, Kemp, MM, Petri Seiler, K, Carrel, HA, Golub, TR, Schreiber, SL, Clemons, PA, Carpenter, AE and Shamji, AF (2013) Multiplex cytological profiling assay to measure diverse cellular states. PLoS One doi: 10.1371/journal.pone.0080999.

12. Wu, PH, Phillip, JM, Khatau, SB, Chen, WC, Stirman, J, Rosseel, S, Tschudi, K, Van Patten, J, Wong, M, Gupta, S, Baras, AS, Leek, JT, Maitra, A and Wirtz, D (2015) Evolution of cellular morpho-phenotypes in cancer metastasis. Sci Rep doi:10.1038/srep18437.

13. Boutros, M, Heigwer, F and Laufer, C (2015) Microscopy-Based High-Content Screening. Cell 163:1314–1325.

14. Lin, JR, Izar, B, Wang, S, Yapp, C, Mei, S, Shah, PM, Santagata, S and Sorger, PK (2018) Highly multiplexed immunofluorescence imaging of human tissues and tumors using t-CyCIF and conventional optical microscopes. eLife 7:e31657.

15. Wawer, MJ, Li, K, Gustafsdottir, SM, Ljosa, V, Bodycombe, NE, Marton, MA, Sokolnicki, KL, Bray, MA, Kemp, MM, Winchester, E, Taylor, B, Grant, GB, Hon, CS, Duvall, JR, Wilson, JA, Bittker, JA, Dančík, V, Narayan, R, Subramanian, A, Winckler, W, Golub, TR, Carpenter, AE, Shamji, AF, Schreiber, SL and Clemons, PA (2014) Toward performance-diverse small-molecule libraries for cell-based phenotypic screening using multiplexed high-dimensional profiling. Proc Natl Acad Sci U S A 111:10911–10916.

16. Wagner, BK and Schreiber, SL (2016) The Power of Sophisticated Phenotypic Screening and Modern Mechanism-of-Action Methods. Cell Chem Biol 23:3–9.

17. Chang, Q, Ornatsky, OI, Siddiqui, I, Loboda, A, Baranov, VI and Hedley, DW (2017) Imaging Mass Cytometry. Cytometry A 91:160–169.

18. Bandura, DR, Baranov, VI, Ornatsky, OI, Antonov, A, Kinach, R, Lou, X, Pavlov, S, Vorobiev, S, Dick, JE and Tanner, SD (2009) Mass Cytometry: Technique for Real Time Single Cell Multitarget Immunoassay Based on Inductively Coupled Plasma Time-of-Flight Mass Spectrometry. Anal Chem 81:6813–6822.

19. Lou, X, Zhang, G, Herrera, I, Kinach, R, Ornatsky, O, Baranov, V, Nitz, M and Winnik, MA (2007) Polymer-based elemental tags for sensitive bioassays. Angew Chem Int Ed Engl 46:6111–6114.

20. Giesen, C, Wang, HA, Schapiro, D, Zivanovic, N, Jacobs, A, Hattendorf, B, Schüffler, PJ, Grolimund, D, Buhmann, JM, Brandt, S, Varga, Z, Wild, PJ, Günther, D and Bodenmiller, B (2014) Highly multiplexed imaging of tumor tissues with subcellular resolution by mass cytometry. Nat Methods 11:417–422.

21. Schapiro, D, Jackson, HW, Raghuraman, S, Fischer, JR, Zanotelli, VRT, Schulz, D, Giesen, C, Catena, R, Varga, Z and Bodenmiller, B (2017) histoCAT: analysis of cell phenotypes and interactions in multiplex image cytometry data. Nat Methods 14:873– 876.

22. Ljosa, V, Caie, PD, Ter Horst, R, Sokolnicki, KL, Jenkins, EL, Daya, S, Roberts, ME, Jones, TR, Singh, S, Genovesio, A, Clemons, PA, Carragher, NO and Carpenter, AE (2013) Comparison of methods for image-based profiling of cellular morphological responses to small-molecule treatment. J Biomol Screen 18:1321–1329.

23. Bougen-Zhukov, N, Loh, SY, Lee, HK and Loo, LH (2017) Large-scale image-based screening and profiling of cellular phenotypes. Cytometry A 91:115–125.

24. Henjes, F, Bender, C, von der Heyde, S, Braun, L, Mannsperger, HA, Schmidt, C, Wiemann, S, Hasmann, M, Aulmann, S, Beissbarth, T and Korf, U (2012) Strong EGFR signaling in cell line models of ERBB2-amplified breast cancer attenuates response towards ERBB2-targeting drugs. Oncogenesis doi:10.1038/oncsis.2012.16.

25. Holliday, DL and Speirs, V (2011) Choosing the right cell line for breast cancer research. Breast Cancer Res 13:215–224.

26. Prat, A, Karginova, O, Parker, JS, Fan, C, He, X, Bixby, L, Harrell, JC, Roman, E, Adamo, B, Troester, M and Perou, CM (2013) Characterization of cell lines derived from breast cancers and normal mammary tissues for the study of the intrinsic molecular subtypes. Breast Cancer Res Treat 142:237–255.

27. Lacroix, M and Leclercq, G (2004) Relevance of breast cancer cell lines as models for breast tumors: an update. Breast Cancer Res Treat 83:249–289.

28. Caie, PD, Walls, RE, Ingleston-Orme, A, Daya, S, Houslay, T, Eagle, R, Roberts, ME and Carragher, NO (2010) High-content phenotypic profiling of drug response signatures across distinct cancer cells. Mol Cancer Ther 9:1913–1926.

29. Bafna, S, Kaur, S and Batra, SK (2010) Membrane-bound mucins: the mechanistic basis for alterations in the growth and survival of cancer cells. Oncogene 29:2893–2904.

30. Kim, SM and Hahn, JH (2008) CD98 activation increases surface expression and clustering of beta1 integrins in MCF-7 cells through FAK/Src- and cytoskeleton-independent mechanisms. Exp Mol Med 40:261–270.

31. Garcia, R, Franklin, RA and McCubrey, JA (2006) EGF induces cell motility and multi-drug resistance gene expression in breast cancer cells. Cell Cycle 23:2820–2826.

32. Blajeski, AL, Phan, VA, Kottke, TJ and Kaufmann, SH (2002) G(1) and G(2) cell-cycle arrest following microtubule depolymerization in human breast cancer cells. J Clin Invest 110:91–99.

33. Ji, J, Zhang, Y, Redon, CE, Reinhold, WC, Chen, AP, Fogli, LK, Holbeck, SL, Parchment, RE, Hollingshead, M, Tomaszewski, JE, Dudon, Q, Pommier, Y, Doroshow, JH and Bonner, WM (2017) Phosphorylated fraction of H2AX as a measurement for DNA damage in cancer cells and potential applications of a novel assay. PLoS One doi: 10.1371/journal.pone.0171582.

34. Molnar, C, Jermyn, IH, Kato, Z, Rahkama, V, Östling, P, Mikkonen, P, Pietiäinen, V and Horvath, P (2016) Accurate Morphology Preserving Segmentation of Overlapping Cells based on Active Contours. Sci Rep doi: 10.1038/srep32412.

35. Sobecki, M, Mrouj, K, Colinge, J, Gerbe, F, Jay, P, Krasinska, L, Dulic, V and Fisher, D (2017) Cell-Cycle Regulation Accounts for Variability in Ki-67 Expression Levels. Cancer Res 10:2722–2734.

36. Van der Maaten, LJP and Hinton, GE (2008) Visualizing High-Dimensional Data Using t-SNE. Journal of Machine Learning Research 9:2579–2605.

37. Jones, TR, Carpenter, AE, Lamprecht, MR, Moffat, J, Silver, SJ, Grenier, JK, Castoreno, AB, Eggert, US, Root, DE, Golland, P and Sabatini, DM (2009) Scoring diverse cellular morphologies in image-based screens with iterative feedback and machine learning. Proc Natl Acad Sci U S A 6:1826–1831.

38. Dao, D, Fraser, AN, Hung, J, Ljosa, V, Singh, S and Carpenter, AE (2016) CellProfiler Analyst: interactive data exploration, analysis and classification of large biological image sets. Bioinformatics 32:3210–3212.

39. Subramanian, A, Narayan, R, Corsello, SM, Peck, DD, Natoli, TE, Lu, X, Gould, J, Davis, JF, Tubelli, AA, Asiedu, JK, Lahr, DL, Hirschman, JE, Liu, Z, Donahue, M, Julian, B, Khan, M, Wadden, D, Smith, IC, Lam, D, Liberzon, A, Toder, C, Bagul, M, Orzechowski, M, Enache, OM, Piccioni, F, Johnson, SA, Lyons, NJ, Berger, AH, Shamji, AF, Brooks, AN, Vrcic, A, Flynn, C, Rosains, J, Takeda, DY, Hu, R, Davison, D, Lamb, J, Ardlie, K, Hogstrom, L, Greenside, P, Gray, NS, Clemons, PA, Silver, S, Wu, X, Zhao, WN, Read-Button, W, Wu, X, Haggarty, SJ, Ronco, LV, Boehm, JS, Schreiber, SL, Doench, JG, Bittker, JA, Root, DE, Wong, B and Golub, TR (2017) A Next Generation Connectivity Map: L1000 Platform and the First 1,000,000 Profiles. Cell 171:1437–1452.

40. Zhou, JX, Isik, Z, Xiao, C, Rubin, I, Kauffman, SA, Schroeder, M and Huang, S (2016) Systematic drug perturbations on cancer cells reveal diverse exit paths from proliferative state. Oncotarget 7:7415–7425.

41. Moffat, JG, Vincent, F, Lee, JA, Eder, J and Prunotto, M (2017) Opportunities and challenges in phenotypic drug discovery: an industry perspective. Nat Rev Drug Discov 16:531–543.

42. Takahashi, C, Au-Yeung, A, Fuh F, Ramirez-Montagut, T, Bolen, C, Mathews, W and O’Gorman, WE (2017) Mass cytometry panel optimization through the designed distribution of signal interference. Cytometry A 91:39–47.

43. Chevrier, S, Crowell, HL, Zanotelli, VRT, Engler, S, Robinson, MD and Bodenmiller, B (2018) Compensation of Signal Spillover in Suspension and Imaging Mass Cytometry. Cell Syst 6:612–620.

44. Caicedo, JC, Cooper, S, Heigwer, F, Warchal, S, Qiu, P, Molnar, C, Vasilevich, AS, Barry, JD, Bansal, HS, Kraus, O, Wawer, M, Paavolainen, L, Herrmann, MD, Rohban, M, Hung, J, Hennig, H, Concannon, J, Smith, I, Clemons, PA, Singh, S, Rees, P, Horvath, P, Linington, RG and Carpenter, AE (2017) Data-analysis strategies for image-based cell profiling. Nat Methods 14:849–863.

45. Ljosa, V, Caie, PD, Ter Horst, R, Sokolnicki, KL, Jenkins, EL and Daya, S (2013) Comparison of methods for image-based profiling of cellular morphological responses to small-molecule treatment. J Biomol Screen 18:1321–1329.

46. Doan, M, Vorobjev, I, Rees, P, Filby, A, Wolkenhauer, O, Goldfeld, AE, Lieberman, J, Barteneva, N, Carpenter, AE and Hennig, H (2018) Diagnostic Potential of Imaging Flow Cytometry. Trends Biotechnol doi: 10.1016/j.tibtech.2017.12.008.

47. Rahmoune, H and Guest, PC (2017) Application of Multiplex Biomarker Approaches to Accelerate Drug Discovery and Development. Methods Mol Biol 1546: 3–17.

48. Wang, L, Yu, Y, Chow, DC, Yan, F, Hsu, CC, Stossi, F, Mancini, MA, Palzkill, T, Liao, L, Zhou, S, Xu, J, Lonard, DM and O’Malley, BW (2015) Characterization of a Steroid Receptor Coactivator Small Molecule Stimulator that Overstimulates Cancer Cells and Leads to Cell Stress and Death Cancer. Cell 28:240–252.

49. Stock, K, Estrada, MF, Vidic, S, Gjerde, K, Rudisch, A, Santo, VE, Barbier, M, Blom, S, Arundkar, SC, Selvam, I, Osswald, A, Stein, Y, Gruenewald, S, Brito, C, van Weerden, W, Rotter, V, Boghaert, E, Oren, M, Sommergruber, W, Chong, Y, de Hoogt, R and Graeser, R (2016) Capturing tumor complexity in vitro: Comparative analysis of 2D and 3D tumor models for drug discovery. Sci Rep doi: 10.1038/srep28951.

50. Chang, Q, Ornatsky, OI, Siddiqui, I, Straus, R, Baranov, VI and Hedley, DW (2016) Biodistribution of cisplatin revealed by imaging mass cytometry identifies extensive collagen binding in tumor and normal tissues. Sci Rep doi: 10.1038/srep36641.

51. Mjos, KD and Orvig, C (2014) Metallodrugs in medicinal inorganic chemistry. Chem Rev 114:4540–4563.

52. Gillet, JP, Varma, S and Gottesman, MM (2013) The Clinical Relevance of Cancer Cell Lines. J Natl Cancer Inst 105:452–458.

53. Osborne, CK, Hamilton, B, Titus, G and Livingston, RB (1980) Epidermal growth factor stimulation of human breast cancer cells in culture. Cancer Res 40:2361–2366.

54. Hau, A, Ceppi, P and Peter, ME (2012) CD95 is part of a let-7/p53/miR-34 regulatory network. PLoS One doi: 10.1371/journal.pone.0049636.

55. Schulz, D, Zanotelli, VRT, Fischer, JR, Schapiro, D, Engler, S, Lun, XK, Jackson, HW and Bodenmiller, B (2018) Simultaneous Multiplexed Imaging of mRNA and Proteins with Subcellular Resolution in Breast Cancer Tissue Samples by Mass Cytometry. Cell Syst 6:25–36.

56. Bray, MA, Vokes, MS and Carpenter, AE (2015) Using CellProfiler for Automatic Identification and Measurement of Biological Objects in Images. Curr Protoc Mol Biol 109:1–13.

57. Swift, ML (1997) GraphPad Prism, Data Analysis, and Scientific Graphing. J Chem Inf Comput Sci 37:411–412.

58. Holten, D (2006) Hierarchical edge bundles: visualization of adjacency relations in hierarchical data. IEEE Trans Vis Comput Graph 12:741–748

59. Angelo, M, Bendall, SC, Finck, R, Hale, MB, Hitzman, C, Borowsky, AD, Levenson, RM, Lowe, JB, Liu, SD, Zhao, S, Natkunam, Y and Nolan, GP (2014) Multiplexed ion beam imaging of human breast tumors. Nat Med 20:436–442.

60. Pomerantz, AK, Sari-Sarraf, F, Grove, KJ, Pedro, L, Rudewicz, PJ, Fathman, JW and Krucker, T (2019) Enabling drug discovery and development through single-cell imaging. Expert Opin Drug Discov 14:115–125.

61. Beghin, A, Kechkar, A, Butler, C, Levet, F, Cabillic, M, Rossier, O, Giannone, G, Galland, R, Choquet, D and Sibarita, JB (2017) Localization-based super-resolution imaging meets high-content screening. Nat Methods 14:1184–1190.

