## Supplementary Informations for "Multidimensional profiling of drug-treated cells by Imaging Mass Cytometry"

**Supporting Figures**

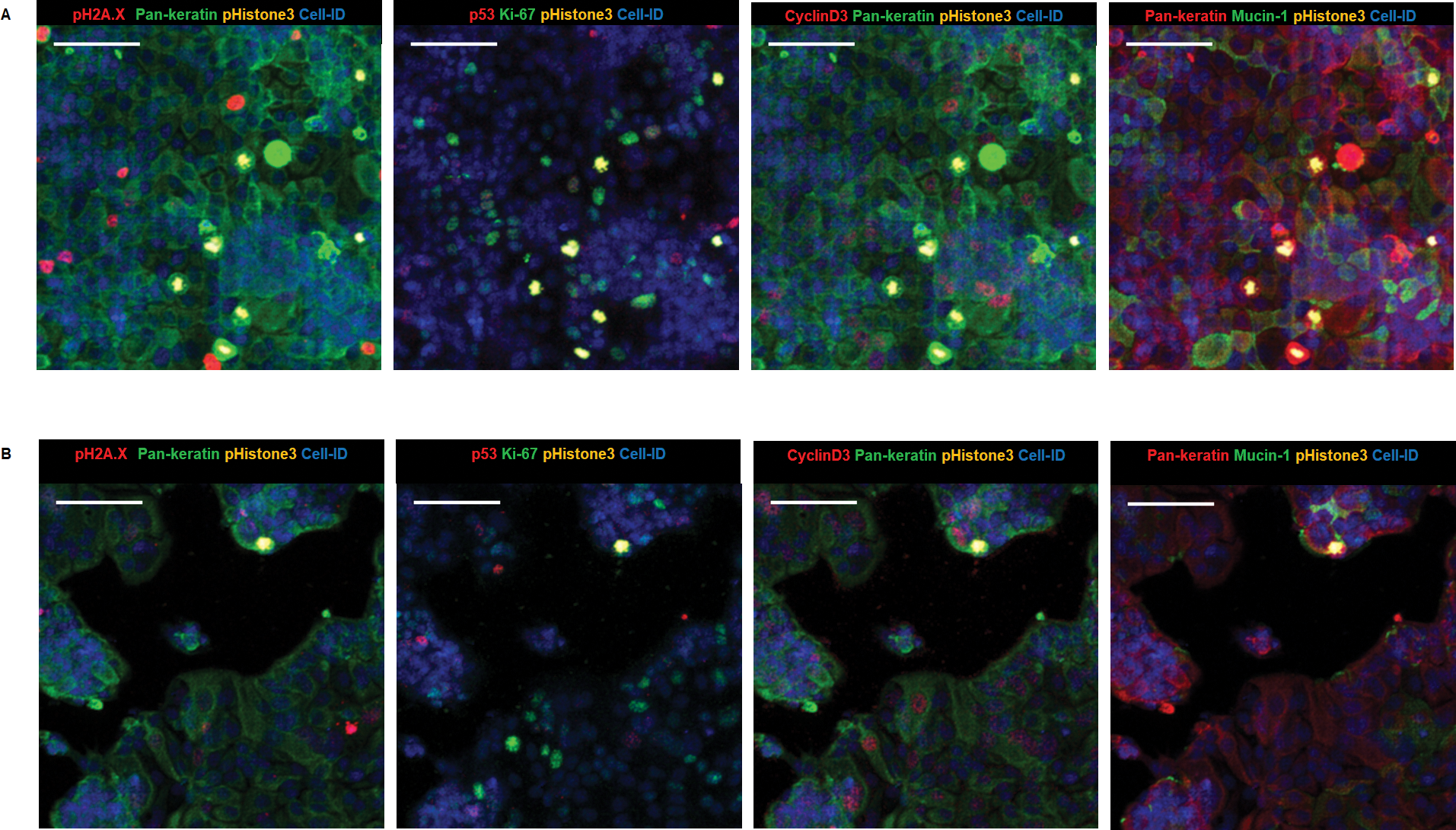

**Figure S1. IMC multiplex image of MCF-7 non-treated and DMSO-treated control cells.**

**Zoom-in colored area of 4-plex non-treated (A) and DMSO-treated control (B) with different combinations of biomarkers, 400 x 400 µm cropped size. Scale bar = 100 µm.**

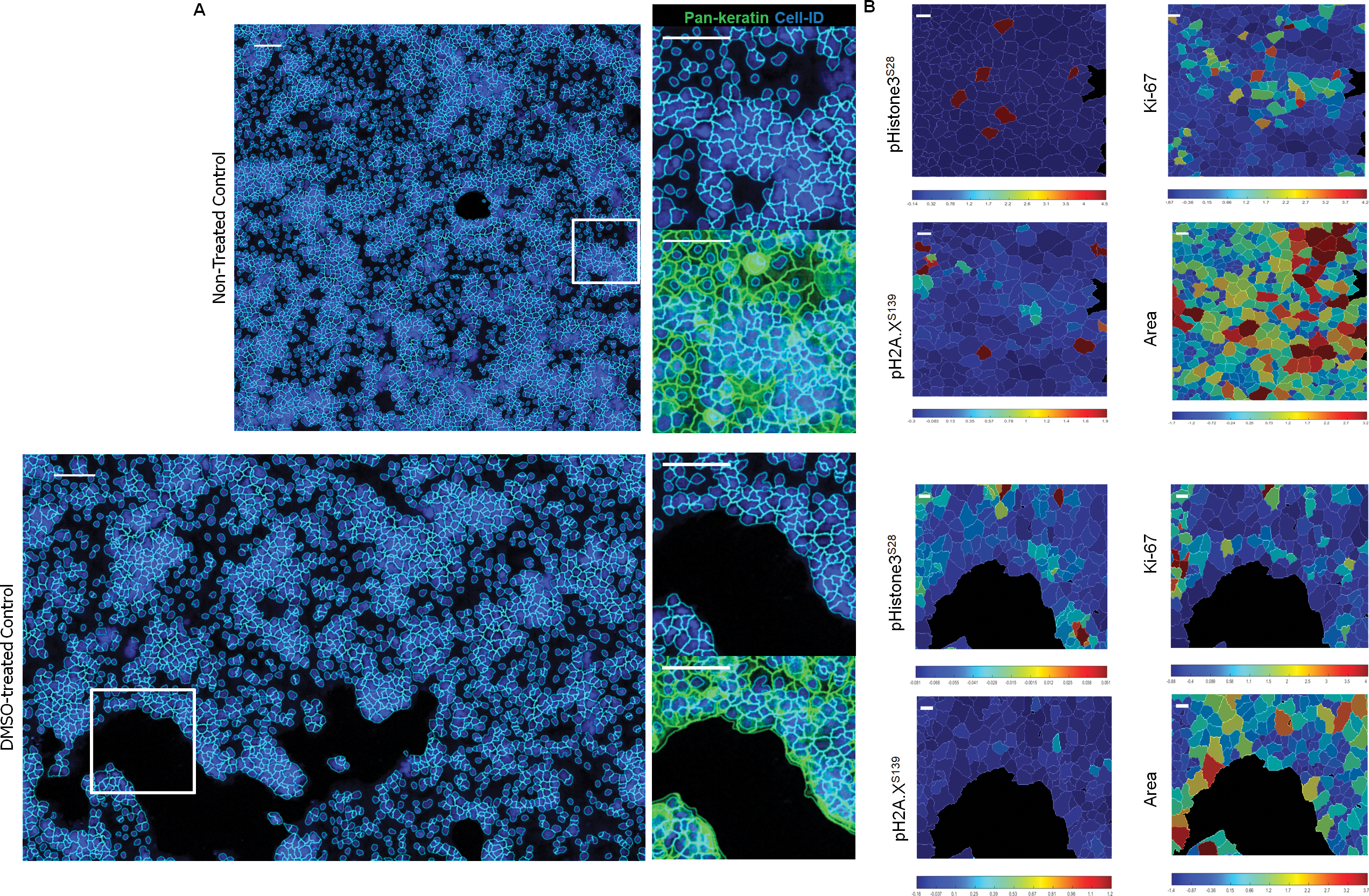

**Figure S2. Cell segmentation and single-marker distribution of MCF-7 non-treated and DMSO-treated controls. (A) CellProfiler overlays of nucleus and cell segmentation masks on color composite images (scale bar = 100 µm). (B) Spatial distribution heat map zoom-in areas of pHistone3^S28^, Ki-67, pH2A.X^S139^ and cell area (scale bar = 10 µm).**

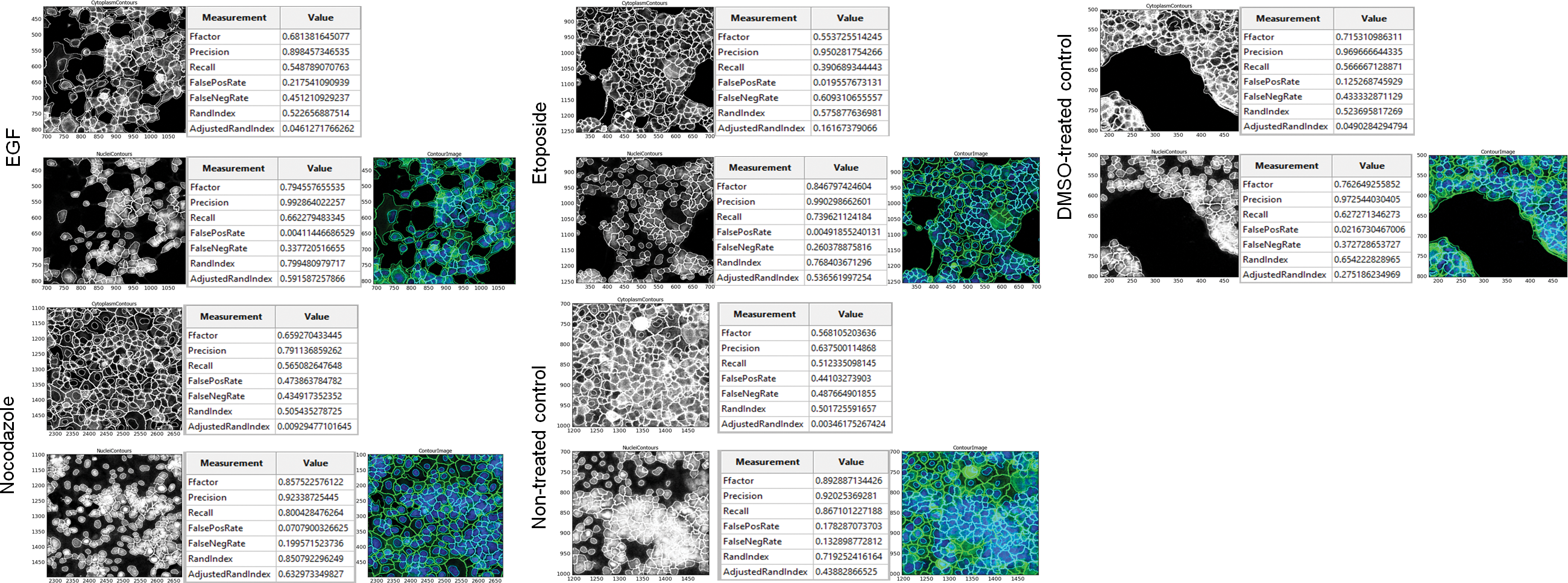
**Figure S3. Segmentation accuracy of nuclei and cytoplasmic contours of identified MCF-7 cells drug-treated using binary image overlap. The identified nucleus and cytoplasm from each cell as object is converted to a binary format called test “image” and overlaid to the binary converted tiff image used for segmentation (DNA-Ir for nuclei identification and pan-keratin for cell identification) as “ground truth”. Both binary images are overlapped to calculate statistics of the closeness from the test image to its ground truth. Performance segmentation statistics parameters is ranging on a scale from 0 to 1, where 1 means perfect overlap and 0 no overlap. Nuclei segmentation shows high precision (number of true positive pixels / (number of true positive pixels + number of false positive pixels) close to 1 over the different images. For cell segmentation the precision is slightly lower but still higher than 0.5 value.**

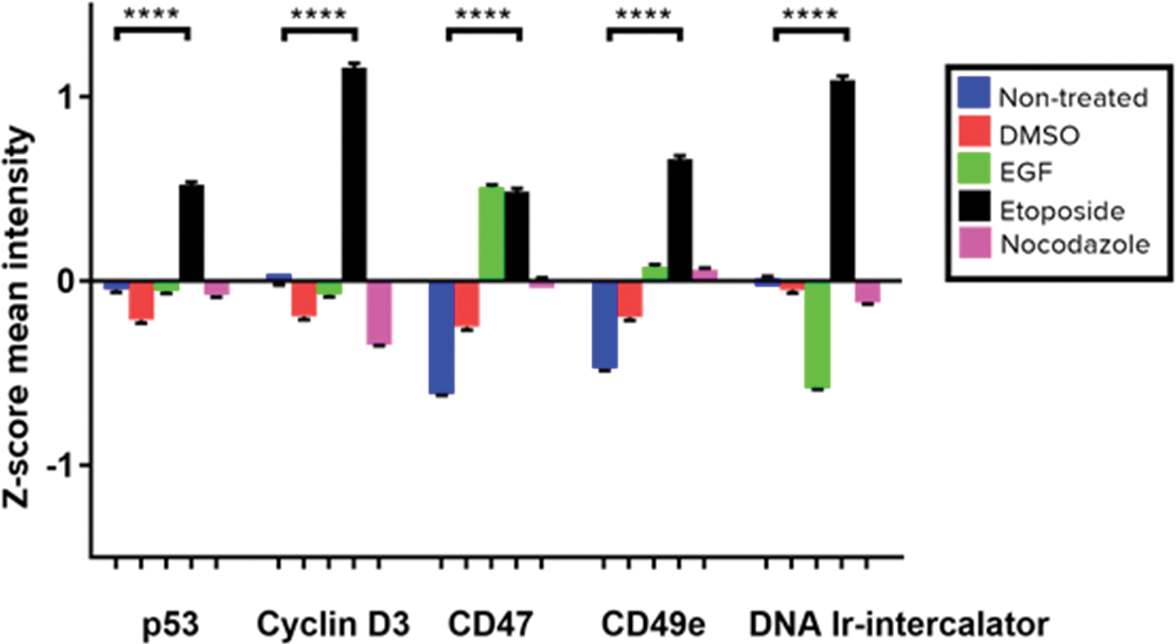

**Figure S4. Comparison of average Z-score expression level of p53, cyclin D3, CD47, CD49e and Cell-ID Intercalator-Ir between each control and drug treatment condition. Data are presented as mean ± SEM. ****P <0.0001 (unpaired t-test).**

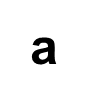

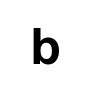

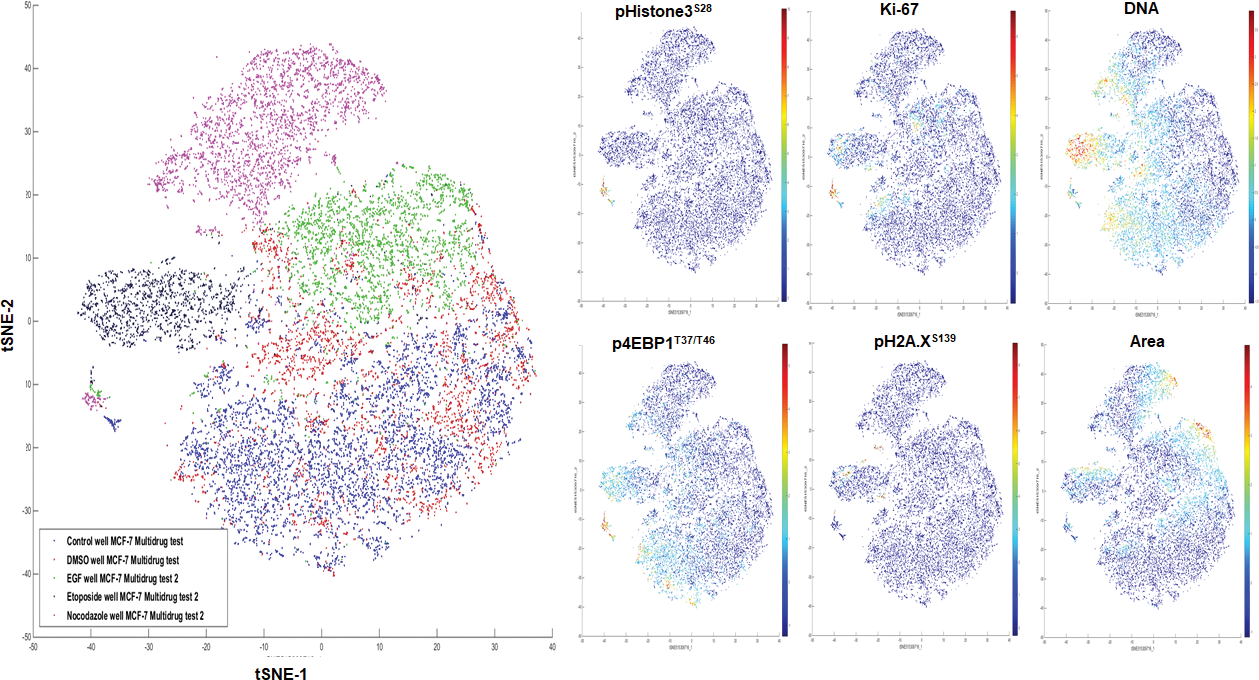

**Figure S5. Unsupervised cluster identification by t-SNE of control and compound-treated MCF-7. (a) Combined colored t-SNE maps of individual cells (EGF, green; etoposide, black; nocodazole, purple), DMSO (red) and non-treated control (blue). (b) t-SNE maps colored according to the expression level of nuclear markers and cell size (Area).**

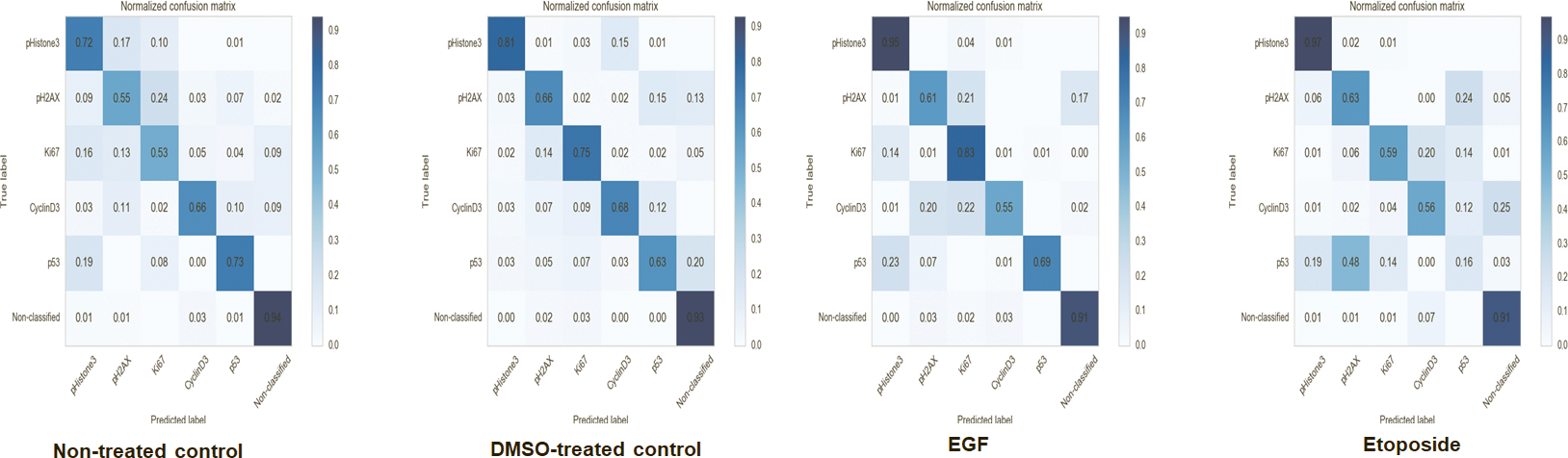

**Figure S6. Fast Gentle Boosting classification matrices of stitched replicates of MCF-7 ROIs per control and drug compound.**

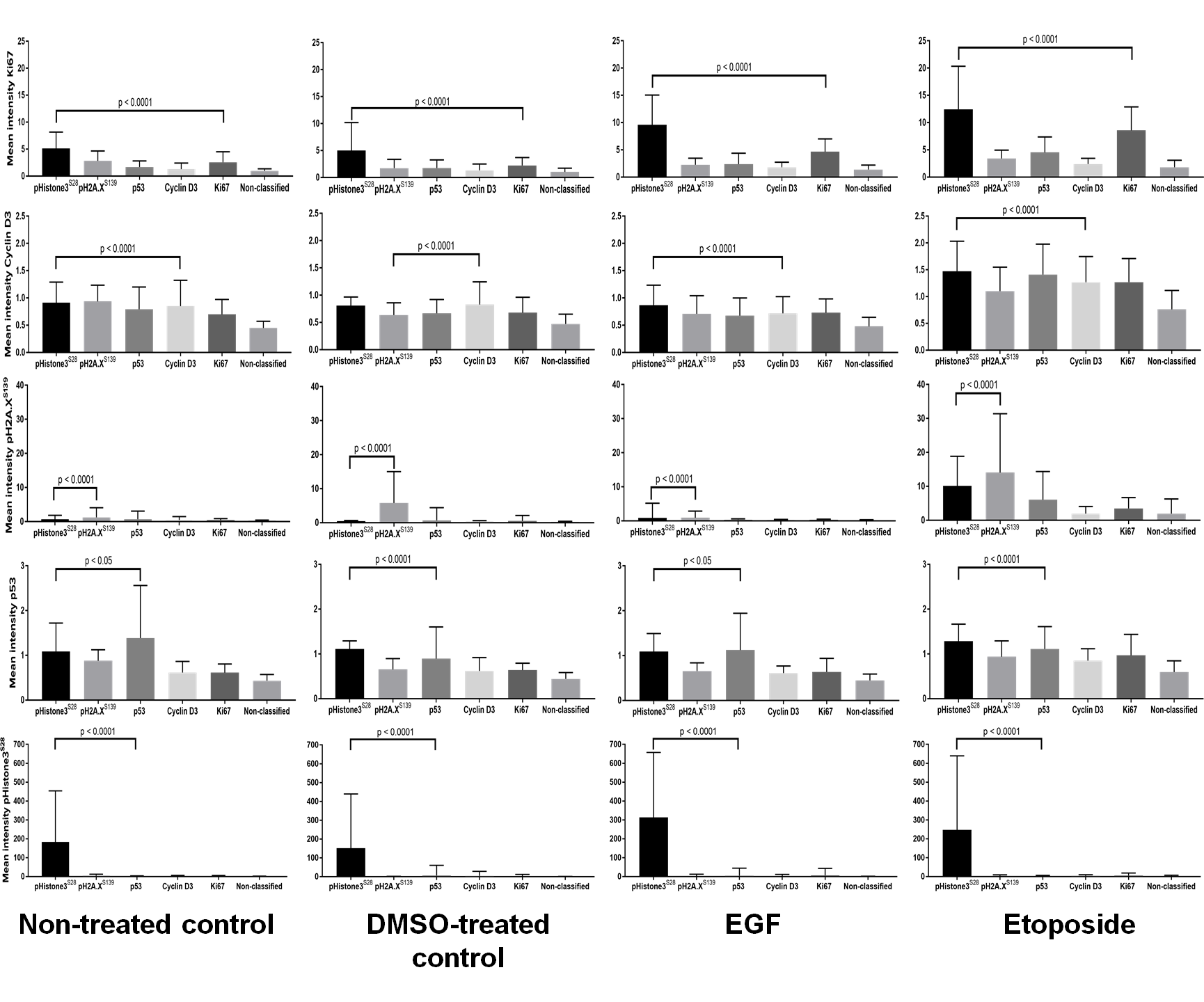

**Figure S7. Comparative expression levels of protein markers Ki-67, cyclin D3, pH2A.X^S139^, p53 and pHistone3^S28^ (rows) per nuclear class and drug treatment condition (column) of MCF-7 cells. Bar charts represent non-normalized mean intensity (+SD) of each nuclear marker to its related class. P < 0.05 and P < 0.0001 (unpaired t-test) are included.**

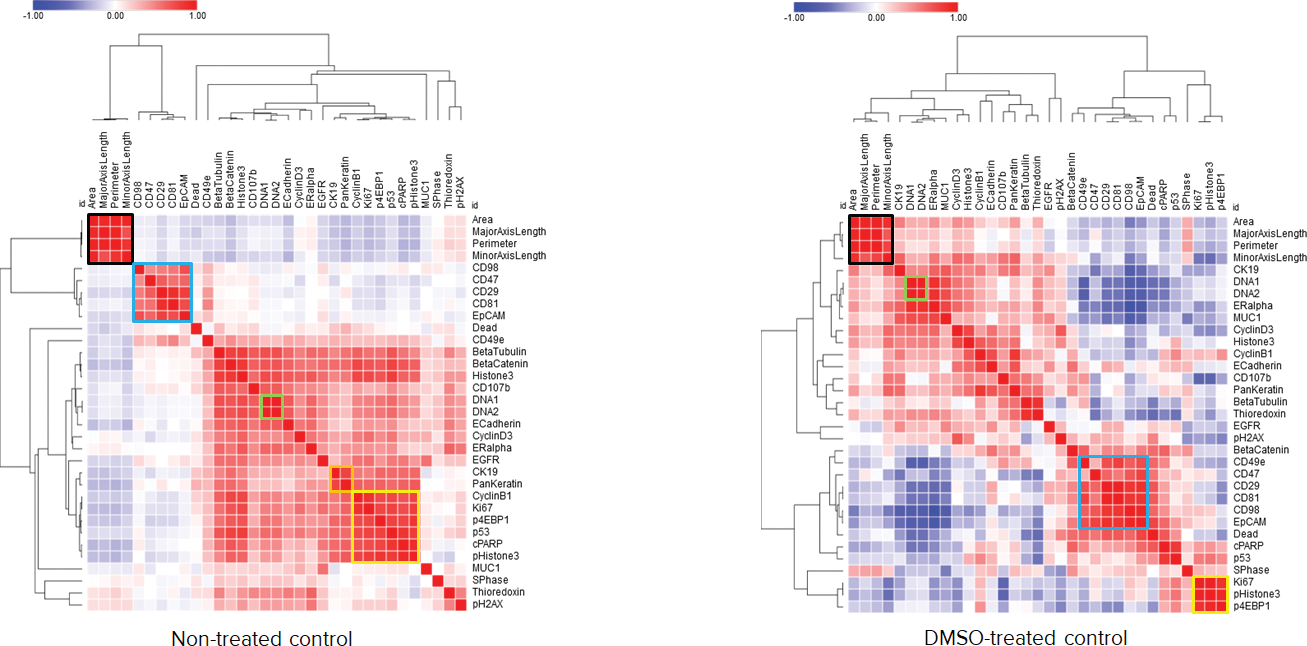

**Figure S8. Heat maps of Pearson correlation coefficients of multiple parameters for non-treated and DMSO-treated controls ROIs on mitotic pHistone3^S28^ cells. Cluster highlighting of cell size parameters (black), surface markers (blue), Cell-ID iridium isotopes (green), pan-keratin and CK19 (orange) and nucleus markers (yellow).**

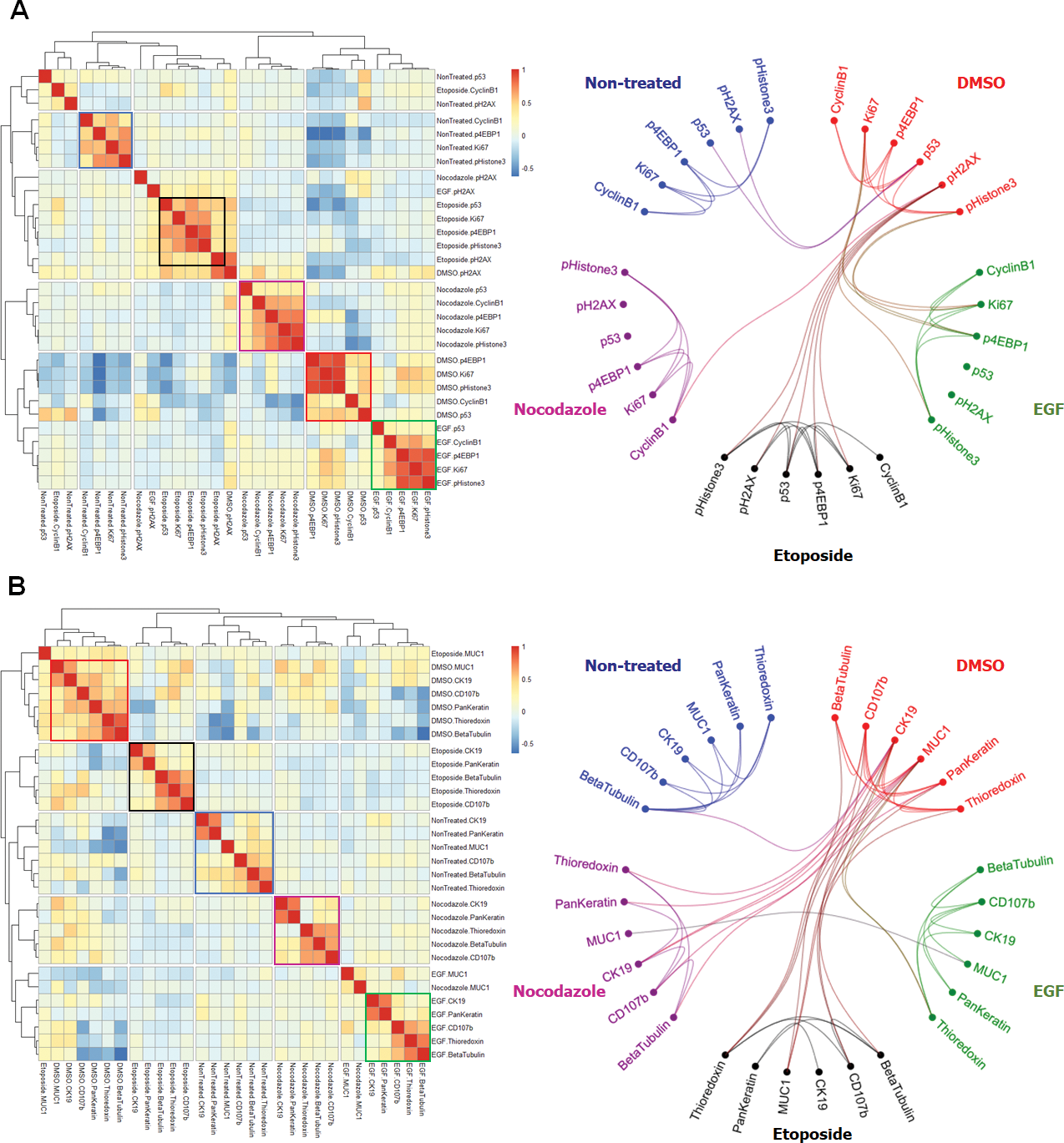

**Figure S9. Graphical representations of Pearson correlation coefficients for nuclei markers (A) and cytoplasmic markers (B) across all controls and drug treatments in the classified population of mitotic pHistone3^S28^ MCF-7 cells. Hierarchical similarities heatmaps show all pairwise correlations values between each protein, with diagonal components highlighted as multiparametric clusters identified for each condition (Non-treated, blue; DMSO, red; EGF, green; etoposide, black; nocodazole, purple). Intra and inter-relationships between these clusters are shown as Hierarchical edgebundle visual graphs. Bundled lines connecting parametric protein nodes correspond to positive correlation values higher than 0.3.**

**Supporting Tables**

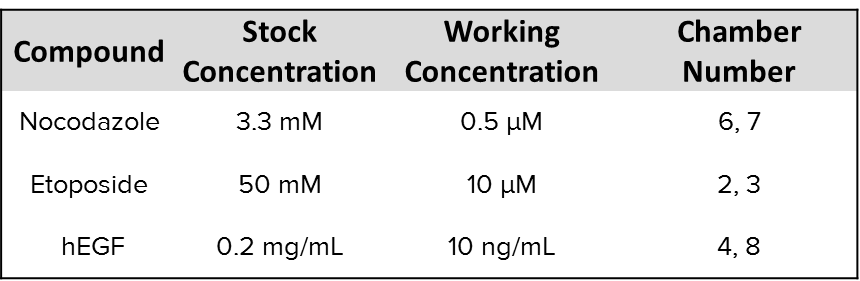

**Table S1. List of chemical compounds used for MCF-7 cell treatment, 48 h exposure.**

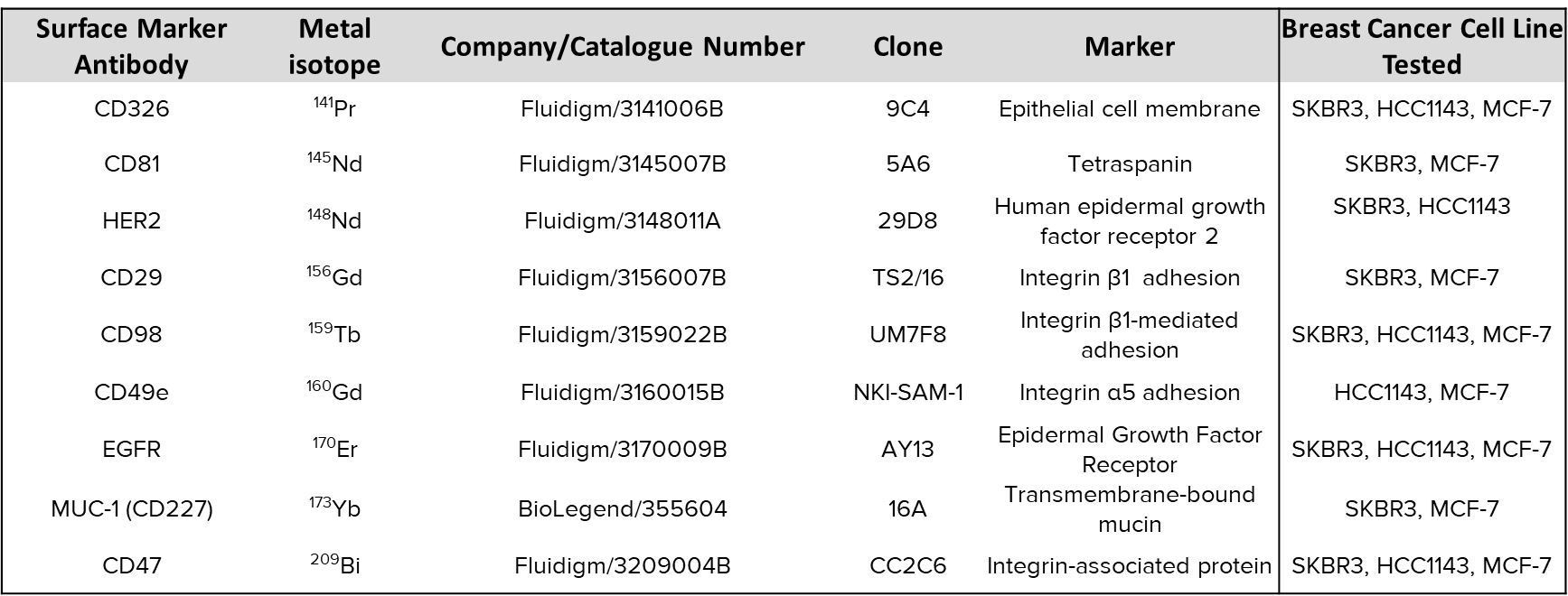

**Table S2. Metal-labeled antibody panel for cell surface markers.**

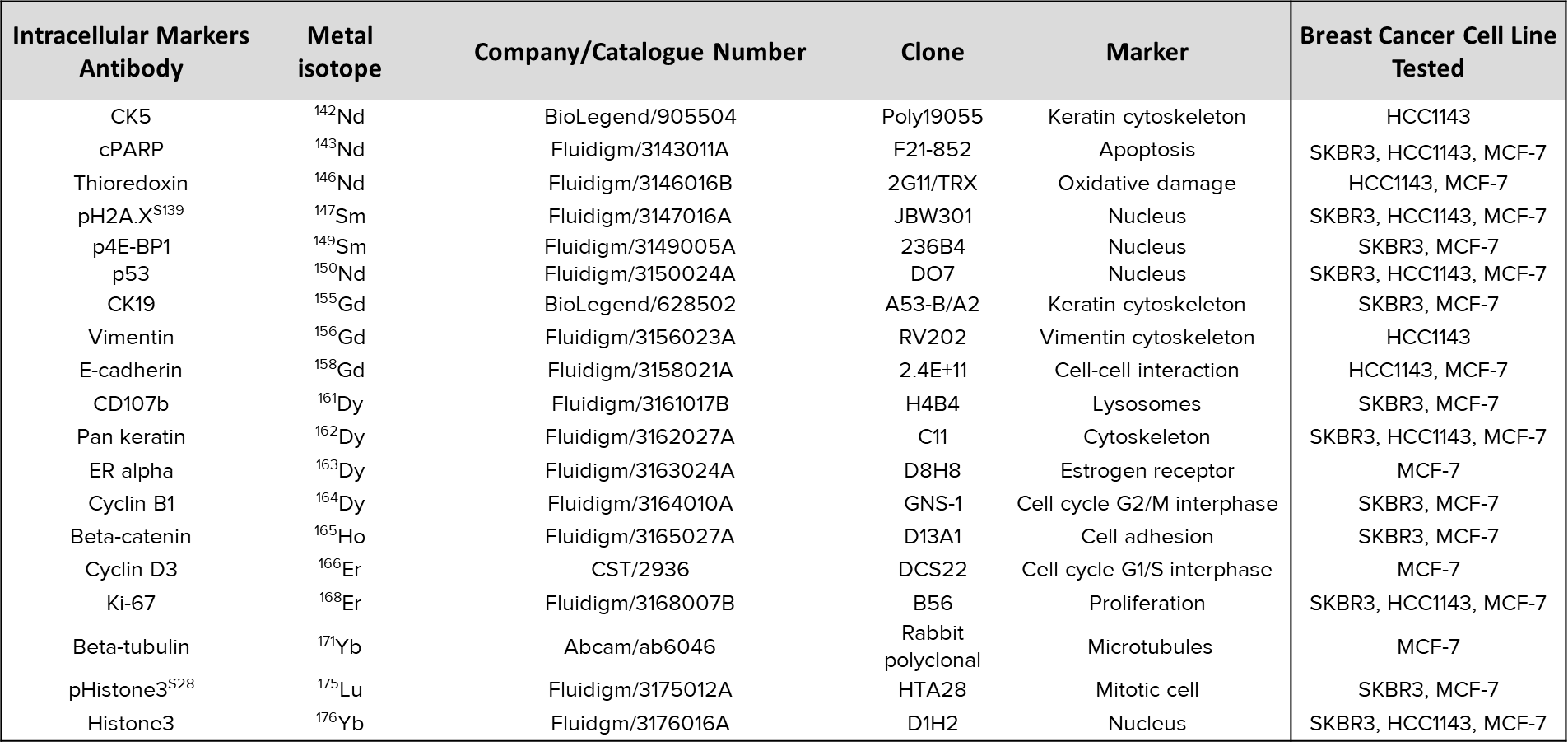

**Table S3. Metal-labeled antibody panel for intracellular markers.**

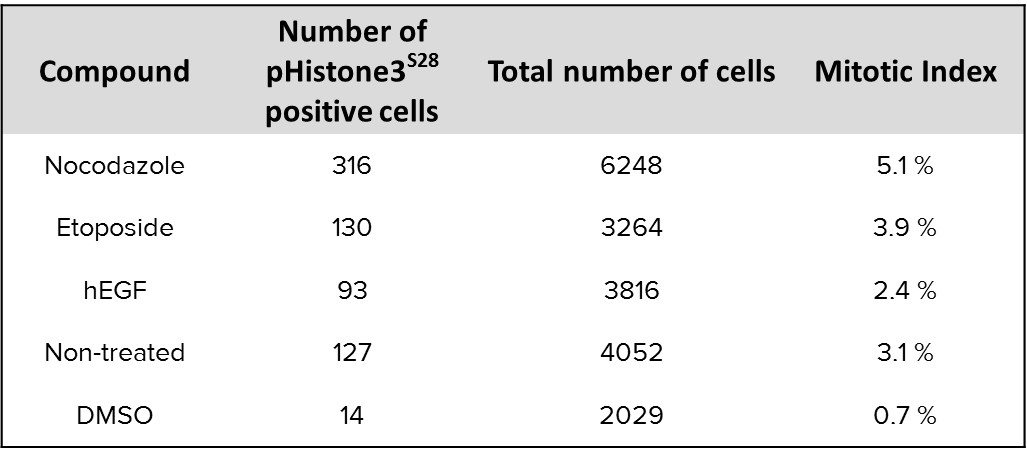

**Table S4. Mitotic index of compound-treated MCF-7 generated by support vector machine classification with Fast Gentle Boosting ruler. Population of pHistone3^S28^ of all ROIs replicates per drug treatment and control were counted and divided by the total number of cells detected to determine the mitotic index of each condition.**
